# SEX-DEPENDENT MODULATION OF WHOLE BRAIN cFOS EXPRESSION BY LIGHT AND MELANOPSIN IN THE MOUSE

**DOI:** 10.64898/2026.09.14.751302

**Authors:** Jacob D. Bhoi, Maya R. Sheth, Lisseth Gonzalez, Andrew Kang, Anna C. Wcislak, Tiffany M. Schmidt

## Abstract

Light is a fundamental feature of an animal’s environment, signaling time of day, weather changes, approaching predators, and other survival cues. The extensive projections from the retina to the brain provide an anatomical substrate for light to adaptively tune circuit function based on changing environmental light. Yet we understand little about the scope and functional impact of light outside of visual and circadian circuits. To address this, we created a whole-brain atlas of light-induced cFos expression in the mouse brain. This approach yielded a strikingly broad pattern of light-driven cFos activation throughout the brain that extended beyond canonical visual circuits, with unexpectedly strong modulation of neuromodulatory centers. These patterns differed starkly in males and females, and were dependent primarily on melanopsin-expressing, intrinsically photosensitive retinal ganglion cells. These findings uncover surprising brainwide patterns of light modulation and are shared as an interactive resource so that they can inspire new, future studies.

**HIGHLIGHTS:**

- Whole-brain atlas of light-induced cFos expression
- Light globally shifts brainwide activity
- Melanopsin regulates baseline neuronal tone in a sex-dependent manner
- Integration of light into neuromodulatory centers could underlie light’s effect

## INTRODUCTION

Light is an inescapable environmental feature that is critical for health and well-being, impacting not only conscious vision, but sleep^1–4^, circadian entrainment^5^, learning^6–9^, mood^10,11^, attention and arousal^12–15^, breathing^16,17^, and many others. Moreover, mistimed light exposure is increasingly recognized as a public health risk and has been empirically linked to numerous negative health outcomes from obesity^18,19^ to anxiety^20^ and even cancer^21–23^. Thus, it is critical to understand the mechanisms through which light influences these myriad behaviors to mitigate negative impacts of aberrant light in an increasingly artificially-lit world and to optimize and uncover new ways that environmental light can be leveraged to improve daily life and health outcomes.

In mammals, light is primarily transduced by three classes of retinal photoreceptors: rods, cones^24^ and the melanopsin-expressing, intrinsically photosensitive retinal ganglion cells (ipRGCs), which are a specialized class of retinal ganglion cell (RGC) that express the photopigment melanopsin (*Opn4*)^25–27^. ipRGCs are unique among RGCs because they integrate the rapid, spatially discrete rod/cone signals with a long-lasting, intrinsic, melanopsin photoresponse driven by environmental luminance levels^28^. In the mouse, ∼40 types of RGC^29–32^ then relay light information from the eye to more than 40 brain regions^33,34^, where they are anatomically poised to broadly impact the brain. Indeed, the ipRGCs themselves comprise the majority of retinal projections outside of canonical visual pathways projecting to more than 20 brain regions throughout the hypothalamus, thalamus, and limbic system^35–37^. ipRGCs have already been linked to a wide range of image forming and non-image forming visual functions^25,28,38,39^, but their widespread projections indicate that they modulate the brain circuits, behavior, and physiology in yet-undiscovered ways.

Despite the anatomical identification of widespread retinal innervation of the brain^33–37^, we still lack a fundamental understanding of the scope and scale of light’s functional impact on the brain. Additionally, the enrichment of ipRGC projections relative to conventional RGCs in non-visual regions makes them strong candidates for modulation of the brainwide function by environmental light. To address these major gaps in a holistic, unbiased way, we developed a quantitative, whole brain atlas of light-induced changes in neuronal activity in the mouse brain using cFos as an indicator of neuronal excitation. We find that acute light exposure causes sex- and melanopsin-dependent changes in brainwide neural activity. Moreover, we find that neuromodulatory centers show significant modulation by light. We have created an interactive website that the field can use as a resource to identify further areas of interest for follow up study that we hope will spawn new and exciting avenues of research. Overall, these results suggest that light does broadly modulate the brain, but that this modulation is gated by melanopsin differentially in male and female animals.

## RESULTS

### Generation of a quantitative, whole brain atlas of light-induced cFos expression

We first sought to identify light-responsive brain regions throughout the whole mouse brain and the contribution of the melanopsin-expressing ipRGCs to these light-dependent changes in activity. To do this, we developed a pipeline that allowed us to generate a quantitative, whole brain atlas of light-induced cFos expression in the mouse brain. We dark-adapted male and female littermate, age-matched control (*Opn4^Cre/+^*) and melanopsin knockout (MKO; *Opn4^Cre/Cre^*) mice overnight and then either exposed them to a 15-minute, broad spectrum, 1000 lux light pulse or kept them in the dark as a control (Figure 1A-B). We delivered the light pulse at CT6, during the deadzone of the photic phase response curve of male and female mice^40,41^, allowing us to better dissociate the circadian and acute effects of light. 90 minutes after light exposure, brains were perfused and processed for whole-brain cFos expression using immunohistochemistry (see Methods). We then used StarDist^42–44^ run through QuPath^45^ to automatically detect cFos^+^ cells, quantified cFos counts in Python, and then determined the location of each individual cFos+ cell by aligning each brain section to the Allen Mouse Brain Atlas Common Coordinate Framework^46^ (CCFv3) using Aligning Big Brains and Atlases^47^ (ABBA) (Figure S1). We then tested whether these automated counts aligned with manual, human quantification by comparing the automated quantification to expert human counts for randomly selected regions throughout the brain (see Methods). Automated counts exceeded expert counts by 21 ± 12 cells per region (mean difference ± SEM; Figure S2). This reflects a higher rate of false positives than false negatives, meaning that automated counts were generally more conservative than human-generated counts (precision, the proportion of detections which are true positives: 0.65 ± 0.06; recall, the proportion of true cells detected: 0.80 ± 0.07). Because detection parameters and imaging conditions were held constant across all animals, this error is expected to affect all groups equally. However, it is important to note that in brain regions with small, dense neurons like the cortex or cerebellar granule layer, the cFos detection undercounts (Figure S3) due to difficulty segmenting individual cFos^+^ nuclei. While this error would impact interregional comparisons, the relative differences across animals within a given region are preserved. Importantly, 87% of cFos^+^ cells were colocalized with the neuronal marker, NeuN (Figure S2), suggesting the majority are neurons.

**Figure 1.**
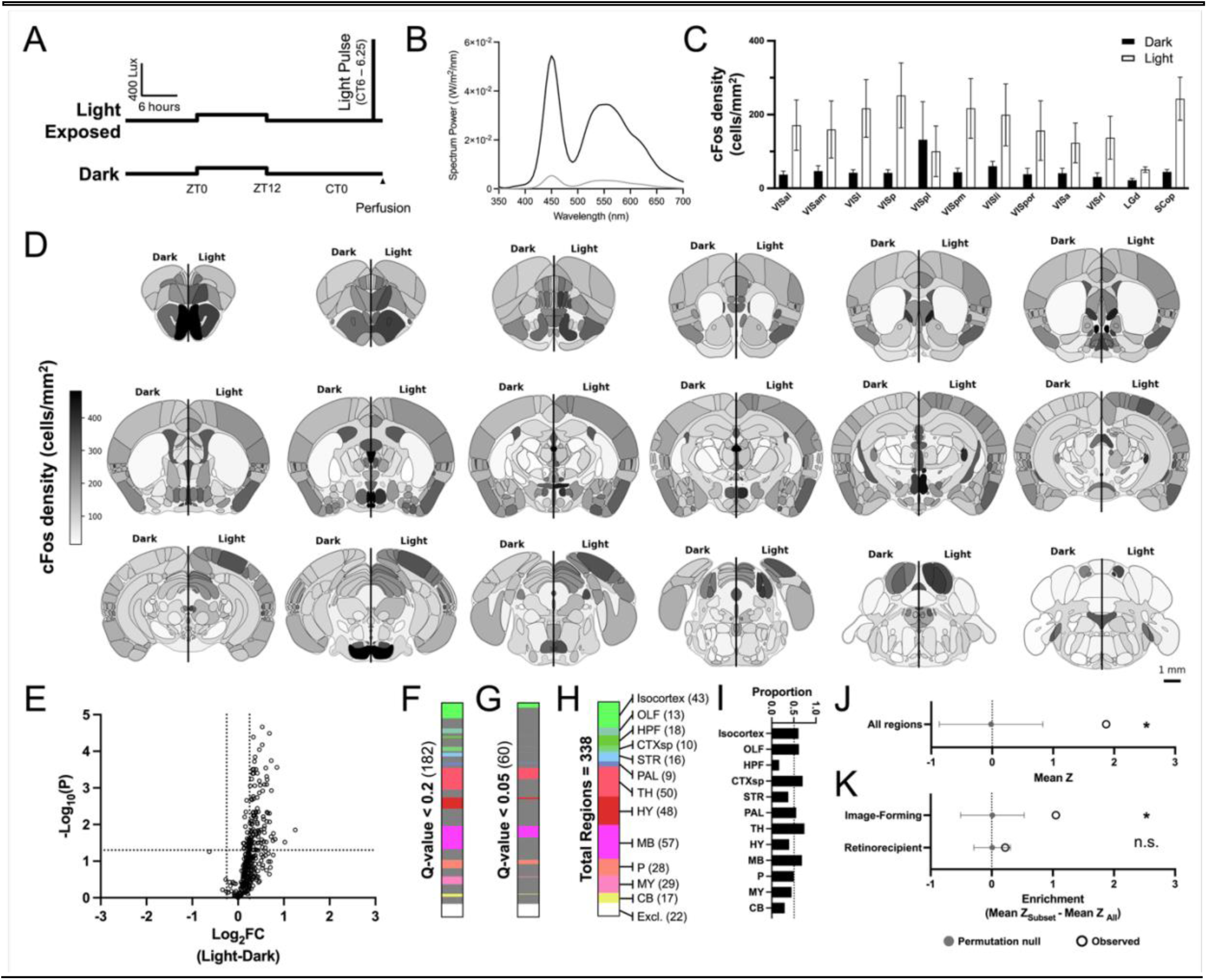
Quantitative, whole-brain atlas of light-induced cFos expression. (A) Schematic of light protocol for dark and light exposed groups, where light-exposed mice receive a 15-minute, 1000 lux pulse of broad spectrum light at CT6. (B) Power along the visible spectrum of light at 100 lux (Grey; normal 12:12 cycle) and 1000 lux (Black; Light pulse). (C) Light-induced cFos expression in the conscious visual pathway of dark-exposed (left, *n* = 11) and light-exposed (right, *n* = 9) control mice. Data analyzed with a repeated-measures mixed-effects model for *light (*p* = 0.048), region, and light x region. Data are Mean ± SEM. (D) Heatmaps of mean cFos density in regions across the brain of all dark-(left) and light-exposed (right) mice. Genotype and sex were pooled. (E) Volcano plot of the Estimate (Log_2_fold-change (FC)) of the main effect of light and the uncorrected -Log_10_(*p*) for each brain region. Positive values indicate higher cFos in light. (F-H) Count of brain regions with a corrected *q*_Light_ < 0.2 (F) and a *q*_Light_ < 0.05 (G) of the total regions tested (H) in the per region linear model. (I) Proportion of regions in each major subdivision of the brain with *q*_Light_ < 0.2. Significance was determined using enrichment analysis (*N* = 12 subdivisions; **MB, *q* = 0.0024, two-sided permutation test with BH correction, *N* = 5000 permutations of animal labels). The line represents an equal distribution of brain regions in each subdivision. (J) Mean *z* of per-region effect of light of observed compared to permuted data (\**p* = 0.029, two-sided permutation test, *N* = 2000 permutations of animal labels). (K) Enrichment analysis for *image-forming (*N* = 13 regions; *p* = 0.040, two-sided permutation test, *N* = 5000 permutations) and retinorecipient regions (*N* = 38 regions; *p* = 0.399, two-sided permutation test, *N* = 5000 permutations). Heatmaps are every 0.5 mm along the anterior/posterior (A/P) axis of the brain. Scale bar = 1 mm.

Prior work has reported that light exposure leads to increased cFos expression throughout the conscious visual pathway^48^. If our pipeline is reliably detecting light-evoked increases in cFos, then we would expect to replicate these findings. To test this, we quantified cFos expression in the conscious visual pathway of male and female controls, including the visual cortex (VIS), dorsolateral geniculate nucleus (LGd), and superior colliculus (SC). In line with previous reports, we measured robust, light-evoked increases in cFos expression in these areas in both male and female control mice, demonstrating that our light stimulus reliably drives detectable increases in cFos expression in these regions (Figure 1C).

### Light induces cFos expression throughout the brain with enrichment in the higher-order visual cortical regions

Given the widespread projections of RGCs to the brain, we next wanted to investigate whether and where light induces cFos expression beyond the conscious visual pathway. To do this, we compared cFos counts of all dark-versus all light-exposed mice. Figure 1D shows heatmaps of cFos density across each grey matter region of the brain. To define which brain regions show light-dependent changes in cFos expression, we fit a linear model on the cFos density of each grey matter region of the mouse brain, with cortical layers collapsed (*N* = 360 candidate regions of which 338 met the minimum of at least 1 mouse per experimental group). Using a false discovery rate (FDR) threshold of *q*_Light_ < 0.2, we found that 182 regions change their cFos expression with the main effect of light (Figure 1F-H), independent of sex and genotype, with nearly all of these regions showing increases in cFos (Figure 1E). Using a more stringent FDR of *q*_Light_ < 0.05, 60 regions showed significant changes with light (Figure 1G-H). Of note, the suprachiasmatic nucleus (SCN) shows similar cFos expression in both light- and dark-exposed mice (*q*_Light_ = 0.96; Supplemental Data File 1), consistent with prior work showing high cFos expression regardless light-exposure at CT6 as well as the lack of light-induced cFos expression after a light pulse delivered during the subjective day^49,50^.

We next compared whether the proportion of significantly different regions is enriched in any of the major subdivisions of the Allen Brain Atlas hierarchy: isocortex, olfactory cortex (OLF), hippocampal formation (HPF), cortical subplate (CTXsp), striatum (STR), pallidum (PAL), thalamus (TH), hypothalamus (HY), midbrain (MB), pons (P), medulla (MY), and cerebellum (CB). To do this, we compared the mean *z*-statistic of each subdivision of regions to the overall population of regions. If light induces cFos expression more highly in specific subdivisions, then we would expect a different mean *z*, with a positive enrichment meaning that subset changed more than the total population. Using this metric, we found that light increased cFos expression in regions across the mouse brain. The thalamus and midbrain showed the largest proportion of significant regions, though only the midbrain showed significant enrichment (Figure 1F-I).

Beyond divisions by anatomical structure, the brain can also be subdivided by function. Decades of research have extensively mapped higher-order visual cortical regions that include the visual cortex (VIS, *N* = 9 regions), retrosplenial cortex (RSP, *N* = 3 regions), and temporal association areas (TEa, *N* = 1 region)^51,52^, and we would expect light to modulate activity in these circuits. Indeed, our analysis revealed that these higher-order visual cortical regions were enriched for light-driven changes in cFos (Figure 1J-K). We also examined whether retinorecipient regions are enriched for light-driven changes in cFos^33,34^. Interestingly, we found that though many individual retinorecipient regions show significant light-dependent changes in cFos, the group as a whole was not significantly enriched (Figure 1J-K). Together, these data suggest that light alone is sufficient to drive cFos expression in many regions of the brain, and higher-order visual cortical regions show enrichment for light’s effect. The fact that 182 regions show significance, but only the MB and higher-order visual cortical regions showed significant enrichment, points to a widespread modulation of the brain by light that extends well beyond canonical visual pathways.

### Brainwide cFos expression is modulated by light and melanopsin in a sex-dependent manner

Recent work has identified previously unknown sex differences in how light and ipRGCs influence behavior^7,13,53–59^. Thus, we next assessed whether acute light exposure alters brainwide cFos expression in a sex- and/or melanopsin-dependent manner. To do this, we fit a linear-mixed model (LMM) with immunohistochemistry batch as a random intercept to measure changes in global cFos expression to look for main effects and interactions between light (dark-versus light-exposed), genotype (control versus MKO), and sex (male versus female) (Figure 2A).

**Figure 2.**
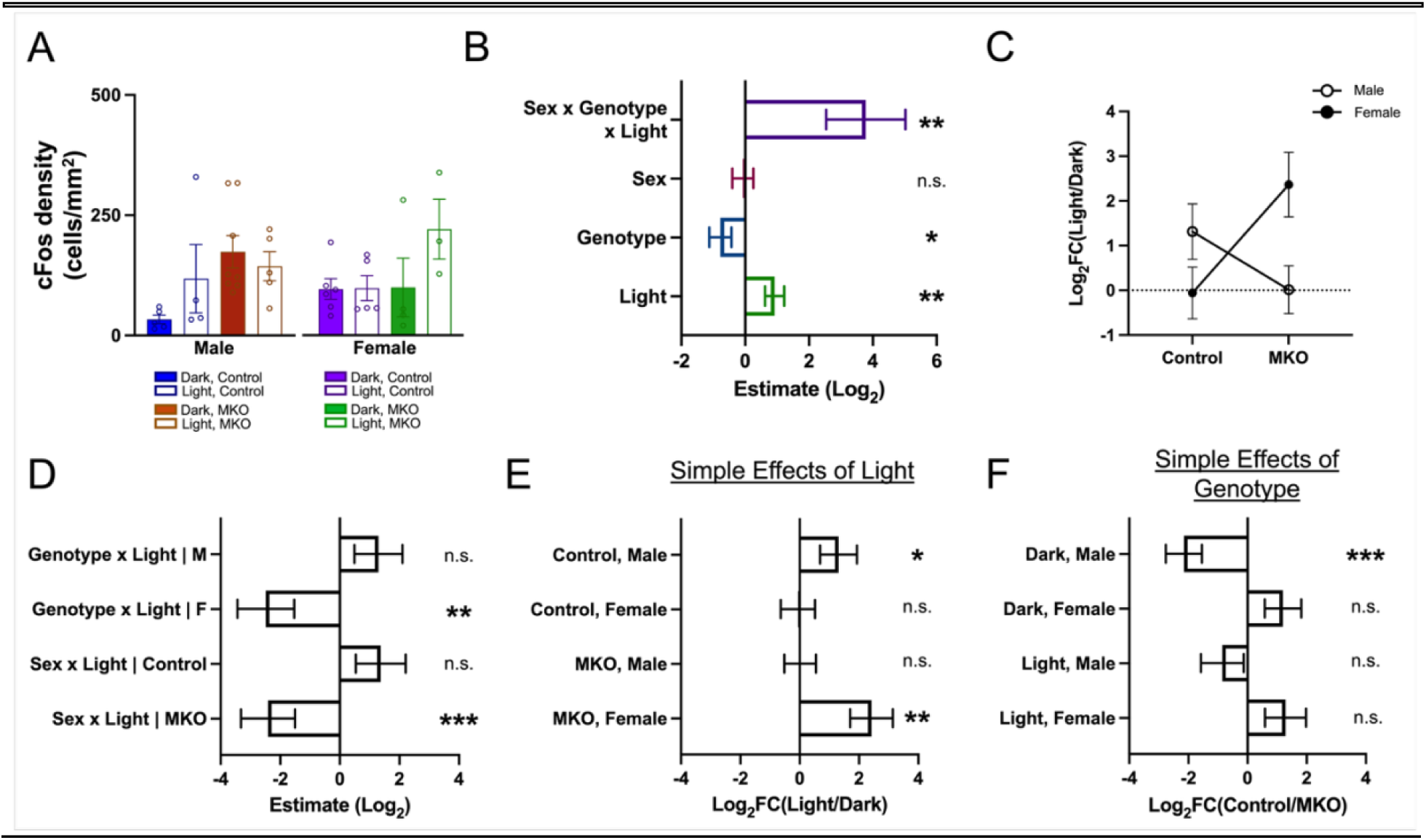
Brainwide influence of light depends on sex and melanopsin. (A) Whole brain cFos density (males: *n* = 4 light controls, 5 dark controls, 5 light MKOs, 8 dark MKOs; females: *n* = 5 light controls, 6 dark controls, 3 light MKOs, 4 dark MKOs). Data are Mean ± SEM. (B) Main effects and three-way interaction. Data are Estimate (Log_2_) ± SE. (C) Log_2_FC(Light/Dark) ± SE in Control and MKO brains of male (open circle) and females (closed circle). (D) Planned decomposition of two-way interactions, computed within each level of the third factor. Each interaction is the difference of one factor’s effect between the two levels of the other. A positive genotype x light interaction in males indicates that light increases cFos more in control than MKO male brains. Data are Estimate (Log_2_) ± SE. (E) Planned decomposition of simple effects of light. Data are Log_2_FC(Light/Dark) ± SE. (F) Planned decomposition of simple effects of genotype. Data are Log_2_FC(Control/MKO) ± SE. n.s. *p* > 0.05; \**p* < 0.05; \*\**p* < 0.01; \*\*\**p* < 0.001. Positive values indicate higher cFos in light, control, and male.

We found significant main effects of light and genotype (Figure S7) but not sex, alongside a significant genotype x sex x light interaction (referred to as GSL-interaction) (Figure 2B and Table S1). This indicates that cFos induction by light differs by genotype in a sex-dependent manner. Furthermore, splitting this interaction by sex shows that the genotype-dependent light effect is opposite in males and females (Figure 2C-D and Table S2). Additional contrasts were measured as planned decompositions. Specifically, control males and MKO females showed significant increases in light-induced cFos induction (Figure 2E), while males showed genotype-dependent changes in cFos expression in the dark (Figure 2F). These results are robust to model start and the systematic exclusion of each sample as well as immunohistochemistry batch, indicating that our findings are not driven by where the model is initialized nor individual experiments or animals (Figure S4). These analyses suggest that there are differential impacts of light exposure based on sex and genotype, which we expand upon further in the sections that follow.

### Light drives a distributed sex- and melanopsin-dependent increase in cFos expression

Given that melanopsin-dependent, brainwide light-induced cFos expression differs between males and females, we next wanted to identify the differences in cFos expression underlying the GSL-interaction. We first visualized regional changes throughout the brain arising from this three-way interaction by splitting the data by sex and plotting the difference in cFos density between the light- and dark-exposed conditions in control and melanopsin knockout (MKO) mice (Figure 3A-B). We also plotted unsubtracted cFos density for controls (Figure S5) and MKOs (Figure S6). These visualizations reveal a striking pattern, showing widespread increases in cFos in the brains of control males that are absent in MKO males (Figure 3A). On the other hand, in control females, we observe little change in cFos expression outside of visual pathways, while MKO females showed widespread light evoked cFos (Figure 3B). This suggests that the genotype-dependent cFos induction by light is distributed throughout the brain, rather than in specific regions, in line with the results of our enrichment analyses.

**Figure 3.**
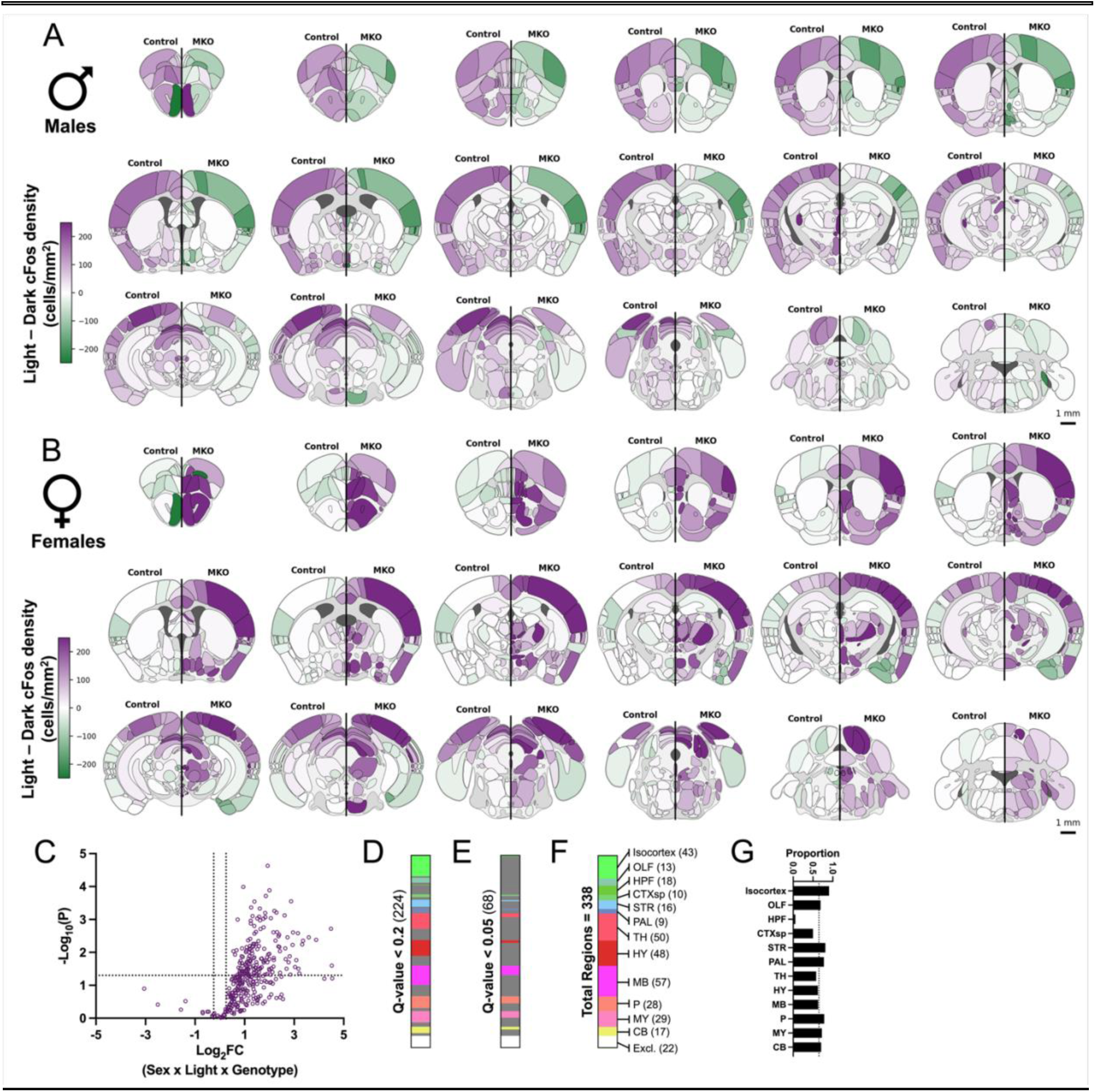
Sex- and melanopsin-dependent light responses are distributed across the brain. (A) Heatmaps of Light - Dark cFos density in male controls (left) and MKOs (right) (mean; *n* = 4 light controls, 5 dark controls, 5 light MKOs, 8 dark MKOs). (B) Heatmaps of Light - Dark cFos density in female controls (left) and MKOs (right) (mean; *n* = 5 light controls, 6 dark controls, 3 light MKOs, 4 dark MKOs). (C) Volcano plot of the Estimate (Log_2_) of the three-way interaction of genotype x sex x light and the uncorrected - Log_10_(*p*) for each brain region. Positive values indicate higher cFos in light, control, and male. (D-F) Count of brain regions with a corrected *q*_GSL_ < 0.2 (D) and a *q*_GSL_ < 0.05 (E) of the total regions tested (F) in the per-region linear model. (G) Proportion of regions in each major subdivision of the brain with *q*_GSL_ < 0.2. Significance was determined using enrichment analysis (*N* = 12 subdivisions; **HPF, *q* = 0.0024, two-sided Freedman-Lane permutation test with BH correction, *N* = 5000 permutations). The line represents an equal distribution of brain regions in each subdivision. Heatmaps are every 0.5 mm along the A/P axis of the brain. Scale bar = 1 mm.

To determine which specific regions show a statistically significant change in a sex- and melanopsin-dependent manner with light, we extracted the GSL-interaction from the per-region linear model as a planned decomposition (Figure 3C). Using a FDR of *q*_GSL_ < 0.2, 224 regions showed significance for the GSL-interaction, indicating that differences in light-evoked cFos expression are widespread, supporting our observation from the density plots (Figure 3D). Using a more stringent FDR of *q*_GSL_ < 0.05, 68 regions showed significance for the interaction, suggesting that even with these more stringent criteria, the impacts of light on the brain are widespread (Figure 3E). We found that no subdivisions showed significant enrichment, suggesting the 224 statistically significant regions are distributed throughout the brain (Figure 3D-G). This suggests that light impacts the brain of control males in a highly distributed, melanopsin-dependent manner. Conversely, we find that light impacts the brain of control females in a much more localized fashion, modulating primarily visual circuits. This specificity appears to be gated by melanopsin signaling from ipRGCs because MKO females show widespread changes in cFos that are similar to those observed in control males.

### Melanopsin sets the baseline neuronal tone in a sex-dependent manner

Our data show widespread increases in light-evoked cFos in male control brains that are absent in MKOs. Surprisingly, females showed the opposite pattern, with MKO females showing widespread light-evoked cFos that was absent in controls (Figure 2E). It is possible that these differences arise from altered baseline cFos in MKO animals. We therefore first visualized this by subtracting control from MKO cFos density in dark-exposed male and female brains to identify genotype-dependent changes in cFos (shown in Figure 4A). The heatmaps reveal that dark-exposed MKO males have widespread, elevated cFos density throughout the brain relative to dark-exposed control males. In contrast, dark-exposed female control brains showed similar cFos density to that of MKOs. This suggests that lack of melanopsin, even in the dark, may drive higher baseline cFos expression in MKO males but not females.

**Figure 4.**
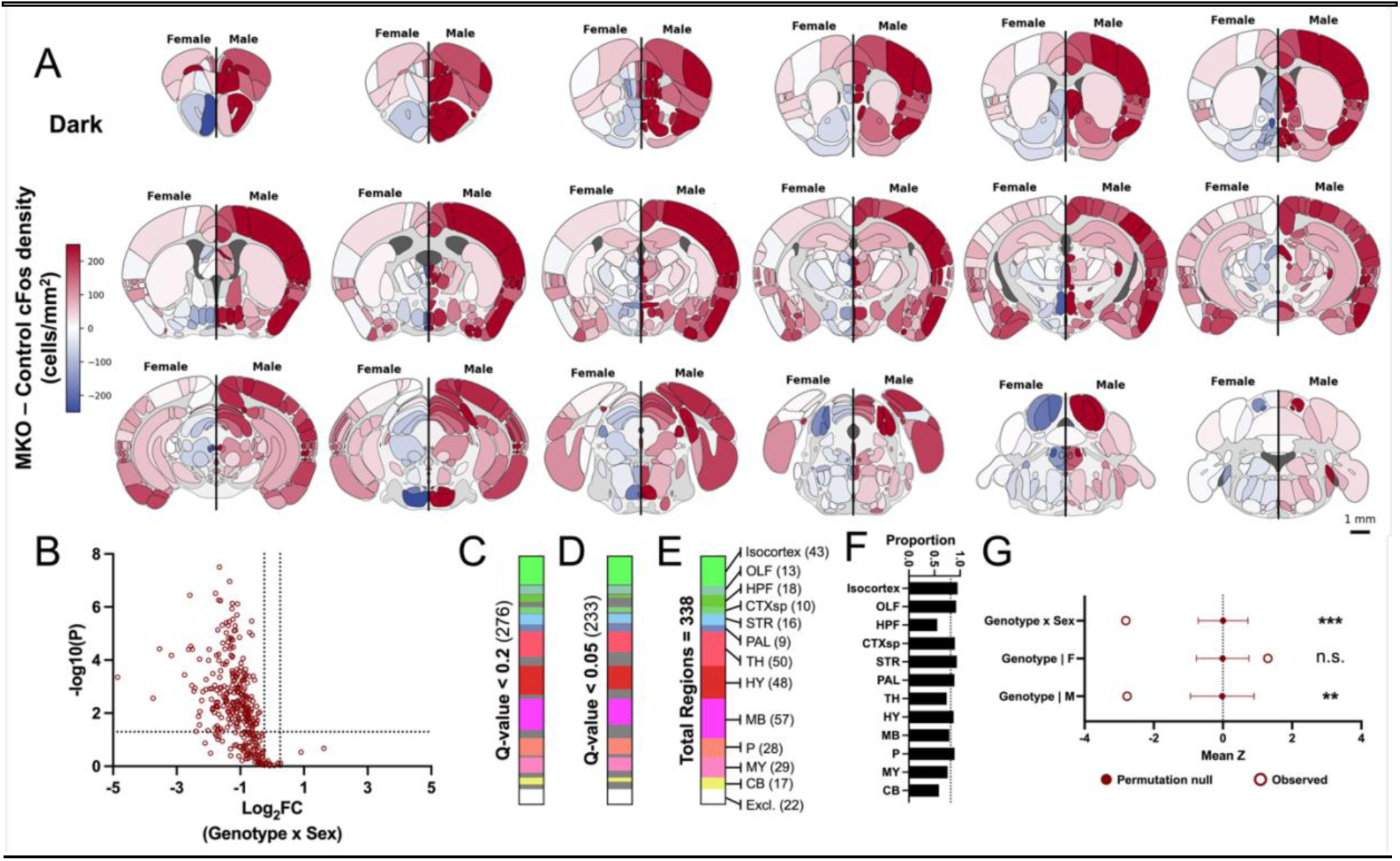
Melanopsin regulates baseline neuronal tone in a sex-dependent manner. (A) Heatmaps of MKO - Control cFos density in dark-exposed females (left) and males (right) (mean; *n* = 6 control females, 4 MKO females, 5 control males, 8 MKO males). (B) Volcano plot of the Log_2_FC of the two-way interaction of genotype x sex in the dark and the uncorrected -Log_10_(*p*) for each brain region. (C-E) Count of brain regions with a corrected *q*_Geno:Sex|Dark_ < 0.2 (C) and a *q*_Geno:Sex|Dark_ < 0.05 (D) of the total regions tested (E) in the per-region linear model. (F) Proportion of regions in each major subdivision of the brain with *q*_Geno:Sex|Dark_ < 0.2. Significance was determined using enrichment analysis (N = 12 subdivisions; *P, *q* = 0.048, two-sided Freedman-Lane permutation test with BH correction, N = 5000 permutations). The line represents an equal distribution of brain regions in each subdivision. (G) Mean *z* of observed compared to permuted data for the per-region two-way interaction of ***genotype x sex in the dark (*p* = 0.001, two-sided Freedman-Lane permutation test, *N* = 2000 permutations) and its decomposition into effects of **genotype in the dark of males (*p* = 0.002, two-sided permutation test; *N* = 2000 permutations of animal labels) and females (*p* = 0.054, two-sided permutation test; *N* = 2000 permutations of animal labels). Heatmaps are every 0.5 mm along the A/P axis of the brain. Scale bar = 1 mm.

To define which brain regions show statistically significant changes, we extracted the two-way interaction of genotype x sex in dark-exposed mice from the per-region linear model (Figure 4B). Using a FDR of *q*_Geno:Sex|Dark_ < 0.2 for discovery, 276 regions showed significance for the genotype x sex interaction in the dark, indicating widespread differences in cFos density (Figure 4C-F). Indeed, even using a more stringent FDR of *q*_Geno:Sex|Dark_ < 0.05, 233 regions showed significance for the interaction, demonstrating that this is a robust effect (Figure 4D). The proportion of significant regions within each subdivision was fairly uniform, with only the pons showing significant enrichment (Figure 4F). To determine how each sex contributes to the genotype x sex interaction in the dark, we compared the mean *z* across all regions to the distribution obtained under *N* = 2000 permutations of animal labels (Figure 4G). The observed mean fell outside the permuted distribution, indicating a genotype x sex effect in darkness beyond that expected by chance. When we look at the simple effect of genotype in males and females in the dark, MKO males exhibit a brainwide increase in cFos expression in dark-exposed conditions compared to control, while dark-exposed female MKOs exhibit no change, with a trend towards a slight decrease in cFos expression after melanopsin loss, further supporting the idea that increased baseline cFos in dark-exposed MKOs is specific to males (Figure 4G). Thus, in males, the primary impact of removing melanopsin is increased baseline cFos in the dark, rather than a failure to increase cFos in the light. This may create a ceiling effect, and could account for the lack of light-evoked cFos increases in MKO males (Figure 3A). However, female MKOs did not show increased baseline cFos in the dark (Figure 4G), despite widespread increases in light-evoked cFos in the female MKO brain (Figure 3B). Thus, in control females, it appears that melanopsin serves to prevent light-evoked cFos expression outside of visual regions, and that this gate is removed in the absence of melanopsin. Overall, these results show that melanopsin has opposing brainwide effects in males and females.

### Light shifts brainwide activity by scaling regional activity

Most behaviors are driven by coordinated changes across brain regions rather than individual brain regions acting in isolation. Thus, we wanted to investigate how the patterns of regional activity underlie the sex- and melanopsin-dependent influence of light on cFos expression. The brainwide effect of light could be produced in two ways. One, light could change which brain regions are active, shifting activity across regions but maintaining a similar overall level of activity. This would present as a change in the spatial patterns of activity. Alternatively, light could change the magnitude of cFos expression in each region, leaving the overall relative patterns of activity intact. This would present as a similar spatial pattern of cFos expression but with changes in its overall amplitude. The per-region linear-model used up to this point determines whether a region changes its cFos expression, but it does take into account whether regions change together, so we next analyzed patterns of cFos expression throughout the population of brain regions using a multivariate approach.

To distinguish between the two possibilities outlined above, we first plotted the log-transformed cFos density of each region in the light compared to in the dark and fit the population with a linear regression. If light increases regional cFos expression amplitude, then we would expect a positive intercept representing the magnitude of the rescaling with the slope indicating whether the scaling is uniform. A slope of one would represent uniform scaling in which baseline activity does not impact light-induced cFos expression. If light recruits particular regions, then individual regions will differ from linear fit, and standard deviation of the residuals (defined as the distance from the fit) should be higher. Finally, if light has no impact, then individual regions would fall along the identity line (slope = 1; intercept = 0), and the goodness of fit for the linear regression would be very tight. Using this approach, we found that control males and MKO females had values that fall above the identity line with a positive intercept (Figure 5A-B), indicating a brainwide increase in cFos in response to light. Additionally, control males and MKO females have a slope less than one, suggesting that the magnitude of light’s effect on cFos expression depends on baseline activity in the dark, with regions that exhibit high baseline cFos showing less induction. In control females the values fall along the identity line (Figure 5A-B), suggesting minimal population influence of light. In MKO males, the slope is less than one, but the intercept does not diverge from zero (Figure 5A-B), consistent with high baseline cFos in the dark of MKO males, suggesting that this group may be approaching a ceiling. However, it is important to note that higher cFos density would be more difficult to detect in cortical or cerebellar regions, as noted above and that the slope for the isocortex does differ from the overall population, indicating that detection capabilities could also contribute to the ceiling effect (Figures S3 and S8). The standard deviation of the residuals is also higher in MKO females (Figure 5B), suggesting that the population mean less accurately captures this group. These data suggest that light acts by shifting population regional activity nonuniformly between genotypes and sexes.

**Figure 5.**
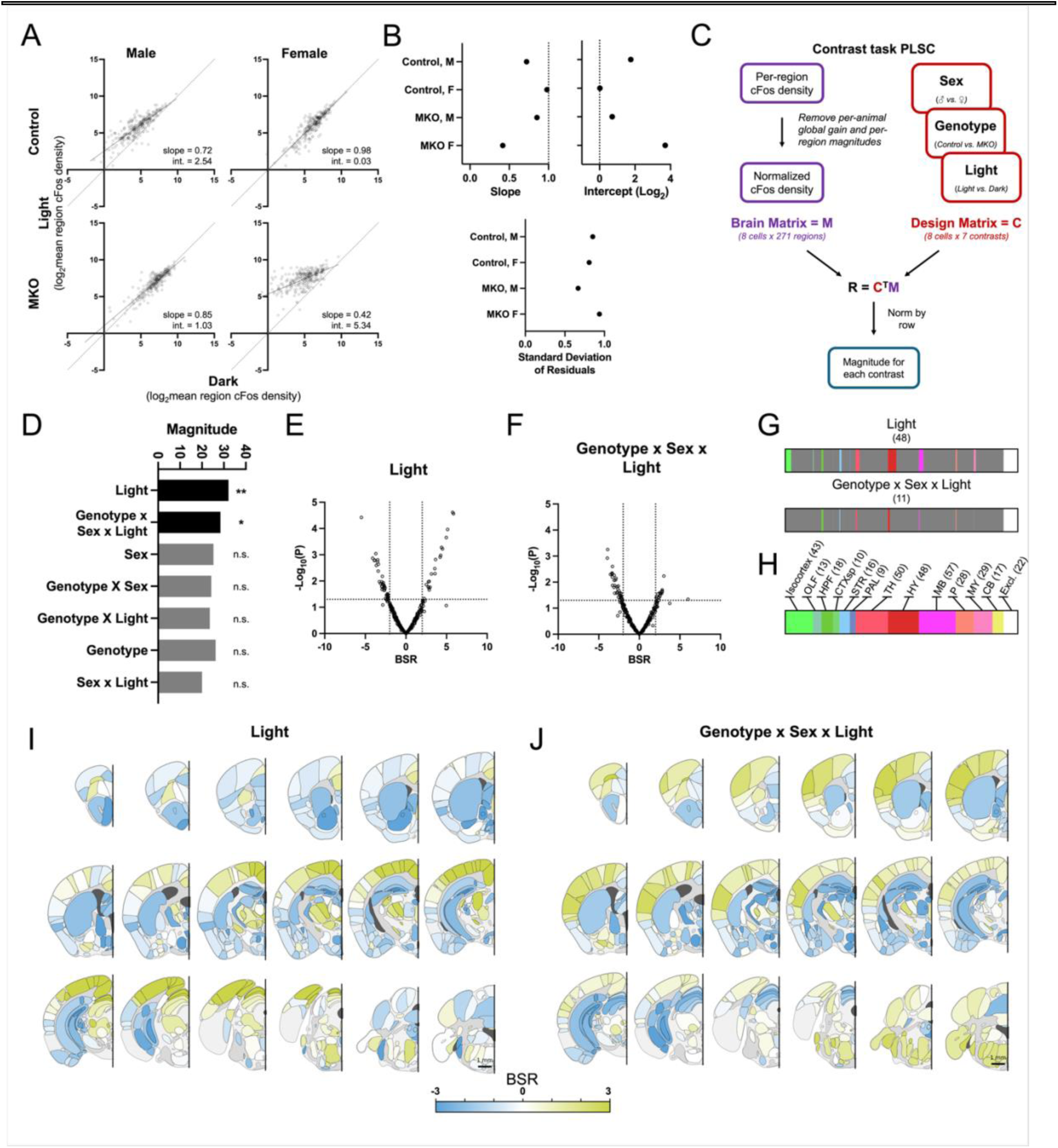
Light shifts the magnitude of brainwide cFos expression rather than recruiting distinct regions. (A) Per-region mean Log_2_ cFos density in the light- vs. dark-exposed brains. Black line is the ordinary least-squares fit. (B) Slope, intercept, and standard deviation of residuals for each fit in (A). Control males **slope (*p* < 0.0005) and **intercept (*p* = 0.0005); control females slope (*p* = 0.33) and intercept (*p* = 0.66); MKO males *slope (*p* = 0.029) and intercept (*p* = 0.096); MKO females **slope (*p* < 0.0005) and **intercept (*p* < 0.0005). *p* values are relative to a slope = 1 and intercept = 0 (dashed lines), the expected outcome if light has no impact. (C) Schematic of contrast task PLSC. (D) Per-contrast magnitudes; **Light (magnitude = 32.0; *p* = 0.0035, one-sided permutation test, *N* = 10000 permutations) and *genotype x sex x light (magnitude = 28.42; *p* = 0.0202, one-sided permutation test, *N* = 9917 permutations) are significant, and all other contrasts are n.s. (E-F) Volcano plot of the bootstrap ratio (BSR) (mean salience / SE) for the effect of light (E) and the genotype x sex x light interaction (F) and the uncorrected -Log_10_(*p*) for each brain region run on mean-subtracted data. (G-H) Count of brain regions with a corrected *q* < 0.2 for the main effect of light (G, top) and the genotype x sex x light interaction (G, bottom) out of the total regions tested (H) in the per-region linear model run on mean-subtracted data. (I-J) Heatmaps of BSR for Light (I) and the three-way interaction of genotype x sex x light (J). Heatmaps are every 0.5 mm along the A/P axis of the brain. Scale bar = 1 mm.

We next wanted to determine which regions show altered cFos beyond the population mean. To do this, we first performed task-centered, partial least squares correlation (PLSC) as described recently by Chiaruttini et al., 2025 in their whole-brain analysis pipeline (Figure S9)^47^. Prior to analysis, regional densities were Log-transformed, the immunohistochemistry-batch contribution was removed from each region, each animal’s brainwide mean was subtracted, and regions were *z*-scored across animals. Subtracting the brainwide mean removes the contribution tested in the global model (Figure 2), and thus, this analysis asks whether there is a pattern across regions once the brainwide shift is removed. Per-region *z*-scoring ensures that individual regions do not dominate the decomposition. Sex, genotype, light, and all interactions were used as main effects with immunohistochemistry batch contributions removed for each region. Because the effect of light and the GSL-interaction are inherently related, this PLSC approach identified these contrasts together, making it difficult to interpret. Thus, we transitioned to a contrast task PLSC (see Methods)^60,61^ to look directly at the contribution of each contrast (Figure 5C). We computed the norm across regions to extract the brainwide salience for each contrast, finding significant effects of light and a significant GSL-interaction (Figure 5D). We next wanted to determine which regions contribute to this pattern of relative activity, we fit a per-region linear-model to the normalized, mean subtracted data for the effects of light (*n* = 48 regions *q* < 0.2; *n* = 19 regions *q* < 0.05) and GSL-interaction (*n* = 11 regions *q* < 0.2; *n* = 0 regions *q* < 0.05) (Figure 5G-H). Together, these data suggest that light and melanopsin have distributed effects and light acts primarily through population-level shifts in gain rather than selective-recruitment, but that a small proportion of regions contribute beyond simple global scaling.

### Neuromodulatory centers show strong, light-dependent modulation

One interesting pattern that emerged from our analysis was that light appeared to have robust sex- and melanopsin-dependent effects on cFos expression in neuromodulatory centers of the brain including the acetylcholine system^62–64^ (magnocellular nucleus, MA; medial septal nucleus, MS; diagonal band nucleus, NDB; substantia innominata, SI; laterodorsal tegmental nucleus, LDT; pedunculopontine nucleus, PPN), the oxytocin system^65^ (supraoptic nucleus, SO; paraventricular hypothalamus, PVH), the dopamine system^66,67^ (substantia nigra pars compacta, SNc; ventral tegmental area, VTA), the serotonin system^68^ (dorsal raphe nucleus, DR; central linear nucleus raphe, CLI), and the norepinephrine system^69^ (locus coeruleus, LC) (Figure 6A). This suggests that environmental light could extend its functional reach through modulation of brainwide activity by tuning, and thus coopting, neuromodulatory circuits to alter activity in their downstream targets.

**Figure 6.**
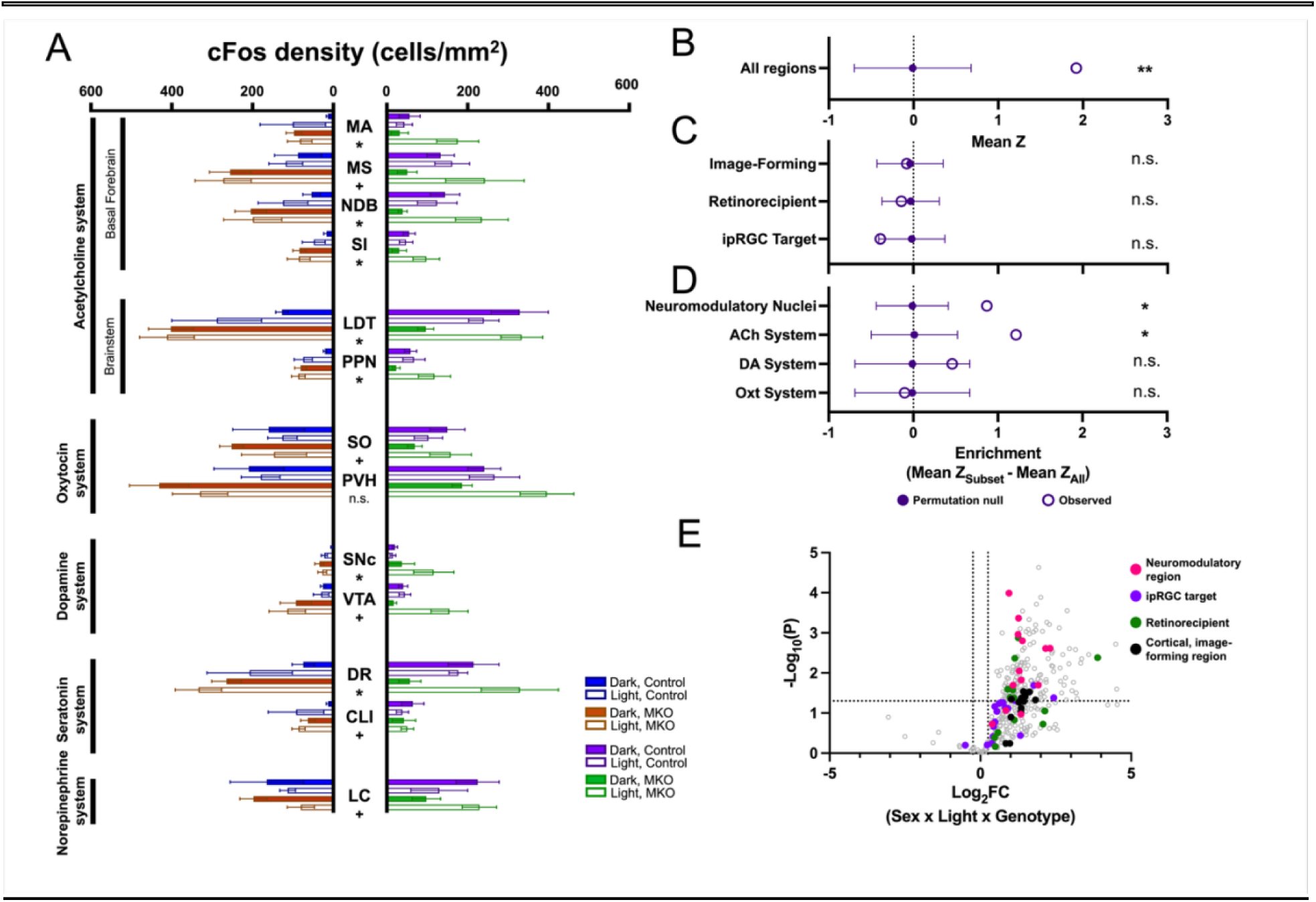
Neuromodulatory centers are among the regions modulated by light. (A) cFos density in neuromodulatory centers in male and females. Significance denotes sex x genotype x light interaction from the per-region linear model. n.s. *q* > 0.2; ^+^*q* < 0.2; \**q* < 0.05; \*\**q* < 0.01; \*\*\**q* < 0.001. Data are Mean ± SEM. (B) Mean *z* of per-region three-way interaction of observed compared to permuted data (\*\**p* = 0.0055, two-sided permutation test, *N* = 2000 permutations). (C) Enrichment analysis for image-forming (*N* = 13 regions tested; *p* = 0.859, two-sided permutation test, *N* = 5000 permutations), retinorecipient regions (*N* = 38 regions tested; *p* = 0.63, two-sided permutation test, *N* = 5000 permutations), and ipRGC targets (*N* = 19 regions tested; *p* = 0.38, two-sided permutation test, *N* = 5000 permutations). (D) Enrichment analysis for all *neuromodulatory nuclei (*N* = 11 regions; *p* = 0.034, two-sided permutation test, *N* = 5000 permutations) as well as the *acetylcholine system (*N* = 6 regions; *p* = 0.021, two-sided permutation test), the dopamine system (*N* = 2 regions; *p* = 0.399, two-sided permutation test, *N* = 5000 permutations), and the oxytocin system (*N* = 2 regions; *p* = 0.866, two-sided permutation test, *N* = 5000 permutations). (E) Volcano plot of the Log_2_FC of the three way interaction of sex x light x genotype and the uncorrected -Log_10_(*p*) for each brain region color-coded based on identity within subsets analyzed for enrichment in (C-D).

We therefore next wanted to determine whether regions housing the neuromodulatory systems are enriched for the GSL-interaction. To do this, we compared the mean *z* of neuromodulatory regions, as well as higher-order visual cortical regions, retinorecipient regions, and ipRGC-recipient regions to determine whether any of these were enriched compared to all of the regions with light-induced cFos expression (Figure 6B-D). Surprisingly, we found that of these groups, only the neuromodulatory centers showed enrichment compared to the overall population (Figure 6C-D). When we analyze each neuromodulatory system individually (acetylcholine, dopamine, and oxytocin), we find that the acetylcholine system is particularly enriched (Figure 6D). Note, the enrichment analysis cannot accommodate missing values (see Methods), and thus, since the serotonin and norepinephrine systems are located in the brainstem, which was occasionally not sampled, thus they could not be included in this analysis. Overall, these results suggest that neuromodulatory centers are particularly impacted by light in a sex- and melanopsin-dependent manner.

### Open-access database of whole-brain light-induced cFos expression

To make this dataset accessible to the community, we developed an open-access, interactive web atlas at [website.name].org (Figure 7).

**Figure 7.**
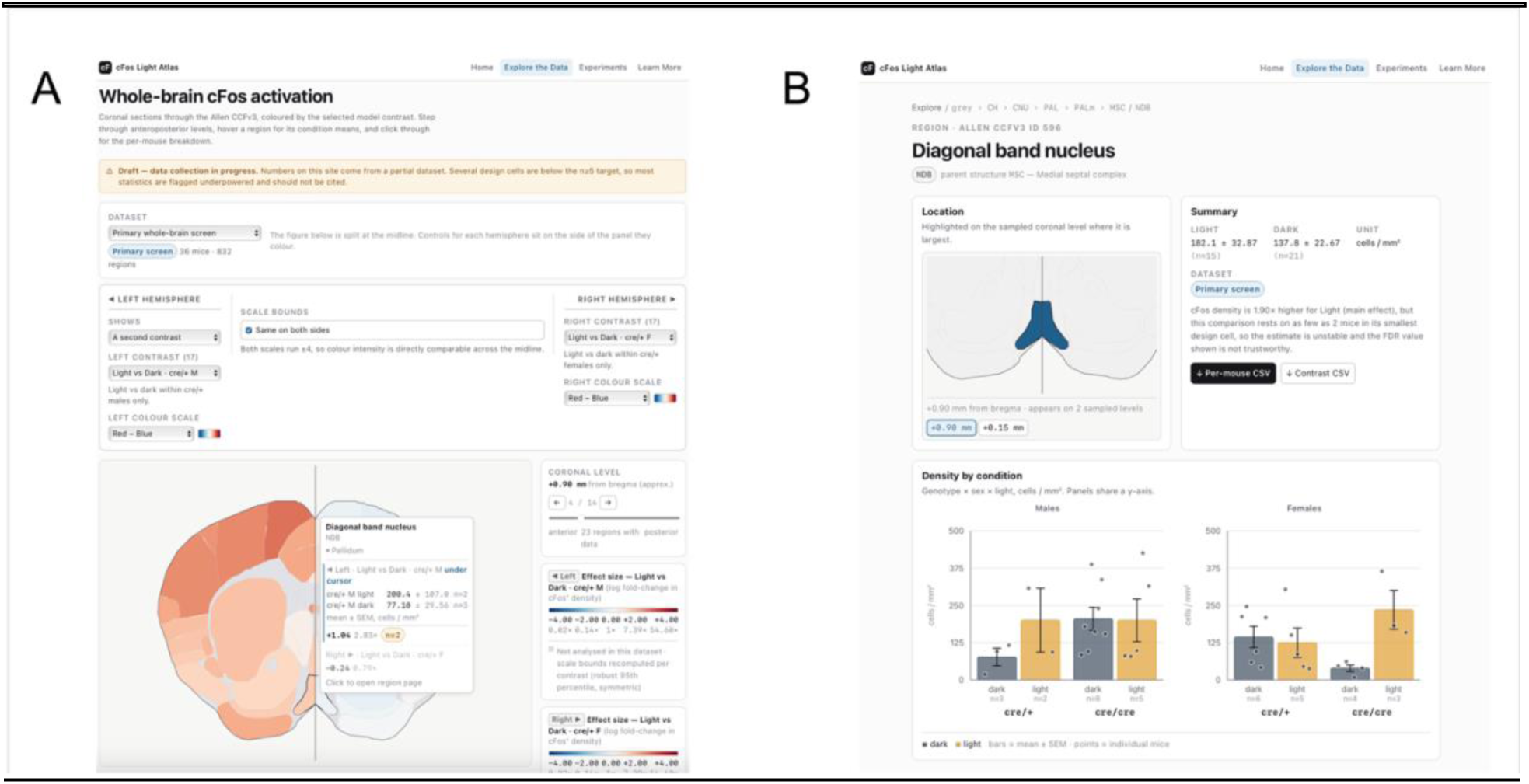
An open-access database of whole-brain light-induced cFos expression. (A) Screenshot of the Atlas “Explore” page of [link]. (B) Screenshot of Atlas “Region” view.

On the Explore page, brainwide results are plotted as coronal heatmaps (Figure 7A). Users can view any comparison of mean cFos densities throughout the brain for each group or the per-region contrast estimates for each main effect (light, genotype, and sex) as well the two- and three-way interactions. The top 25 regions are displayed with their estimate and SEM, ranked by the per-region linear-model *q*-value. Hovering over a region on the coronal heatmaps makes that region’s name and estimate ± SEM appear, and clicking opens a per-region page with individual data points plotted for each group, and the values (estimate ± SEM; *p*) for each contrast (Figure 7B). The experiments page documents the study design, sample sizes, and analysis parameters for each dataset in the atlas database (currently *N* = 1). The website also links to the raw data and analysis scripts for download.

This site is built to expand as more experiments are incorporated measuring other features of how light influences brainwide cFos and other immediate early gene expression throughout the brain. This will allow comparative work across datasets. Ultimately, we hope this resource will support researchers to make testable predictions about how light influences downstream neural circuits and behaviors.

## DISCUSSION

Here, we generated a quantitative, whole brain atlas of light-induced cFos expression in male and female, control and MKO mice. We find that just 15 minutes of light exposure, during a time of day that has no impact on the circadian clock^40,41^, broadly impacts neuronal activity throughout the brain. This whole-brain approach identified hundreds of light-modulated brain regions and showed that the impacts of light are not confined merely to retinorecipient targets, but extend beyond those targets to alter activity throughout the brain in a sex- and melanopsin-dependent manner. Unexpectedly, we found stark differences in how melanopsin signaling shapes light’s impact on the male and female brain.

Our data suggest light scales brainwide cFos expression rather than reorganizing which regions express cFos, because regional changes tend to scale together (Figure 5A). This type of multiplicative scaling is a feature of gain modulation at the single-cell level^70^ suggesting sex- and melanopsin-dependent effect of light may act through gain modulation at the circuit level. However, it is important to note that cFos density does not have the resolution to distinguish between changes in individual neuronal sensitivity (i.e. gain) versus recruitment of additional neurons in each region. Thus, one benefit of this whole brain approach is that it identifies regions for future studies using more precise approaches in intact brains to further distinguish these possibilities.

To date, the majority of studies measuring light-evoked changes in cFos have done so through the lens of circadian dysregulation and/or have targeted cFos expression in a few, specific regions^10,49,71–77^. For example, one study chemogenetically stimulated ipRGCs during the subjective night, finding that this induced cFos in regions of the hypothalamus and amygdala^71^. Excitingly, our study at a different circadian time and using light instead of chemogenetics, also identified 5 of the 7 regions as significant for the GSL-interaction (*q*_GSL_ < 0.2; Supplemental Data File 1). This demonstrates that light has acute, widespread effects beyond impacting the circadian system and that findings with our automated pipeline are replicable using more targeted approaches.

It is notable that light widely affects the brain and that neuromodulatory regions were enriched for sex- and melanopsing-dependent effects on light-induced cFos expression. Neuromodulatory systems project diffusely throughout the brain and are thus anatomically poised to alter circuit function on a brainwide scale based on changes in environmental lighting conditions. Additionally, neuromodulatory systems operate on the timescale of minutes to seconds, driving brainwide changes in activity akin to the population scaling of per-region cFos expression we observe (Figure 5A). The enrichment of neuromodulatory regions, and specifically the acetylcholine system, in the GSL-interaction (Figure 6) is consistent with a model in which light is integrated and signaled via the brain’s neuromodulatory architecture to acutely impact brainwide neuronal activity. In line with this model, RGCs send direct input to the acetylcholine system, with ipRGC input to one of its nuclei the SI^36^, as well as the oxytocin system, with M1 ipRGCs providing input to the SO^78^. This provides a neuroanatomical substrate for light to be integrated into these systems. In fact, recent work demonstrated that dopamine release in the lateral nucleus accumbens acutely tracks environmental luminance, and the effect of light persisted in MKO animals^79^. Future studies should use similar approaches to directly investigate whether and how light is integrated into these neuromodulatory systems, particularly the acetylcholine system (both the basal forebrain and brainstem systems) as well as the serotonin and norepinephrine systems.

Our data support an emerging picture that light has fundamentally different roles in modulating male and female neuronal activity. Our findings show opposing impacts of melanopsin loss in males and females, with melanopsin tuning baseline activity in a sex dependent manner (Figures 3-4). Recent work has demonstrated that females have higher melanopsin expression, and this differentially impacts circadian and acute systems^53^. Female mice show a larger photic phase response than male mice, and this sex difference is abolished in MKOs. In contrast, control male and female mice show similar acute, negative masking responses, but a female-specific deficit is unmasked in MKOs^53^. In a study of a newly described, associative learning in mice called long-term threat avoidance (LTTA), control male and female mice show similar LTTA behavior, but lack of melanopsin again unmasks a sex difference, with males showing decreased LTTA and females showing increased LTTA^7,54^. In males, this is due to melanopsin tuning the baseline activity of the perihabenula, a direct ipRGC target^7^. This baseline tuning of the perihabenula by melanopsin in males is a focal observation of the brainwide effect this dataset supports. Together, these data reveal a pattern in which MKO males and females have diverging phenotypes. These male/female differences have important implications for understanding differences in male and female health outcomes. For example, human studies have also shown that sex differences^13,55,56^, with differential health outcomes from shift work^80,81^ (i.e. mistimed light) and in higher susceptibility of females to seasonal affective disorder^82,83^ (i.e. too little light). Sex differences in how light affects health and behavior may arise from sex differences in melanopsin-established baseline and how that influences the impact of acute changes in environmental light. Our study highlights just how far-reaching the impacts of acute light exposure truly are, and how much we have yet to uncover about the behavioral and physiological consequences of light and how those consequences differ in males and females.

Recent work by Komal et al., 2026 suggests a model in which there is a gate at the ipRGC to SCN connection^84^, explaining the deadzone in the circadian phase response curve and the insensitivity of the SCN to cFos induction from light during the day^49,73^. Komal et al propose that this gate is driven by changes in population sensitivity to depolarization block between day and night. The threshold for depolarization block is a feature that varies amongst ipRGC subtypes^85^ as well as within the M1 ipRGC population to set cellular-level sensitivity to different ranges of environmental luminance^86^. Our data support this model, showing no light-dependent cFos induction in the SCN (*q*_Light_ = 0.96; Figure 3A and Supplemental Data File 1). Our data suggest that in control females, this gate extends beyond the SCN whereas in control males it may be restricted to the SCN. This could be due to sex differences in depolarization block threshold of M1 populations, potentially due to higher melanopsin expression in females^53^ shifting their population sensitivity to be more sensitive, though our cFos atlas alone cannot answer this. Thus, different population sensitivity could result in a more or less widespread gate. Additionally, our data suggest that in melanopsin null mice, the gate is alleviated, increasing male MKO baseline cFos expression throughout the brain, potentially limiting light’s ability to further increase cFos expression from an already high baseline, whereas in female MKOs, alleviating the gate permits widespread cFos induction by light (Figure 4 and Figure 5).

This atlas will serve as a resource for research investigating the influence of light on a wide range of neural circuits, and our hope is that this resource will spawn multiple new avenues of research and grow as new experiments are added. Researchers can use the interactive website to query specific brain regions to generate testable hypotheses related to how light influences specific behaviors and neural circuits. Furthermore, as brainwide atlases become more widespread, for example measuring cFos across circadian time^87^ or between behaviors^47^, they can be used comparatively to disentangle the acute effects of light from other aspects of subconscious visual processing and behaviors. Ultimately, this will feed into future research characterizing the complex and sexually dimorphic circuits underlying the impacts of light on behavior and will help humans adapt to a world with increasing exposure to artificial light.

### Limitations of the study

One limitation of the study is the use of cFos as a proxy for neuronal activity. cFos is a valuable tool for defining changes in neuronal activity, but it has some important limitations relevant to the interpretation of these data including: brain regions and cell types show differences in cFos induction, neuronal activity can vary in many ways of which cFos is only a binary signal, cFos can only detect only positive changes in neuronal activity, and cFos has poor temporal resolution^88–93^.

Additionally, we only looked at changes in the expression of cFos, one example of an immediate early gene. Recent work has shown that different immediate early genes are differentially induced by neuronal activity in various populations^47,94–96^, and the intracellular pathways that lead to immediate early gene expression differ amongst populations^92^. Thus, it is possible that the use of different immediate early genes could reveal further differences.

Another potential limitation of our study is the potential impact of any developmental effects due to the loss of melanopsin. In fact, there is robust evidence for melanopsin’s role in development^97–101^, including impacts on oxytocin release^8^. However, we think developmental effects are unlikely to fully explain our results for two reasons. First, many of the acute impacts of light on behavior are only minimally impacted in MKOs^2^, and the effects of melanopsin loss can be rescued by chemogenetically activating ipRGCs, suggesting development in the absence of melanopsin does not dramatically alter the underlying circuitry^7,35,102^. Second, the opposing impact of MKO in males and females means that if our results were due to a compensatory mechanism, it must be both extensive and opposite in males and females, which seems unlikely. Future work performing a quantitative, whole-brain analysis after chemogenetic activation or inhibition of ipRGCs as well as melanopsin knockout in adulthood would help dissociate any developmental effects.

## METHODS

### Animals

All procedures were approved by the Animal Care and Use Committee at Northwestern University (Protocol number: IS00003845). Animals were housed in a vivarium under 12:12 light/dark (LD) cycle conditions with 100-400 lux light during the light phase, depending on rack position, and *ad libitum* access to food and water. The temperature ranged from 21 to 23°C, and the humidity ranged from 30% to 70%. Adult (P60-P90) male and female littermate mice *Opn4^Cre/+^* and *Opn4^Cre/Cre^* mice^103^ (RRID: IMSR_JAX:035925) with a mixed B6/129 background were used for all experiments. Mouse sex was determined by external anatomy.

Mice were anesthetized with an intraperitoneal (IP) injection of Avertin. Thoracotomy and perfusion were used as secondary methods of euthanasia following deep anesthesia. All efforts were made to minimize pain and discomfort throughout the experiments.

### Light pulse

One day prior to the light pulse, mice were transferred to individual cages and placed in a light control box with a 12:12 LD cycle aligned to the vivarium housing with light intensity of 100 lux during the light phase. Then, the day of the light pulse, the lights remain off from circadian time (CT) 0 CT0 - CT6 to dark adapt the mice. Then, there is a 15-minute, broad-spectrum pulse of 1000 lux light from CT6 - CT6.25, during the deadzone of the circadian phase response curve^40,41^. After the light pulse, mice remain in the dark until they are euthanized (CT7.5 - CT8). Dark-exposed control mice remain in darkness from CT0 until they are euthanized on the day of the light pulse (CT7 - 8.5).

### Immunohistochemical (IHC) procedures and microscopy

Light and dark-exposed mice and genotypes were balanced across perfusion sessions. In order to constrain perfusions to precise CT, perfusion rounds were typically either male or female mice. Males and females of both genotypes and light conditions were pooled for IHC to minimize the impact of batch effects.

90-minutes after the light pulse, mice were anesthetized by IP injection of Avertin and were perfused with PBS followed by 4% PFA in PBS. Brains were dissected and fixed for 24 h at 4°C in 4% PFA in PBS. Brains were then cryopreserved in 30% sucrose in PBS at 4°C until their density increased to that of the solution (2 - 3 days), and then they were embedded and flash frozen in OCT using a −80°C freezer. Tissue was stored at −20 °C for up to 2 weeks prior to sectioning.

Coronal, 60 μm sections were obtained on a Leica CM1950 cryostat. Tissue sections were collected into 1X PBS and stored for up to 1 week at 4°C prior to immunohistochemistry (IHC). Every third slice was collected and processed, meaning the atlas samples 60 μm out of every 180 μm of tissue along the anterior/posterior (A/P) axis.

Sections were washed and then blocked at 4°C overnight in PBS with 6% normal donkey serum and 0.3% Triton-X prior to incubating in primary antibody solution for 2-3 nights at 4°C (Table 1). We used two monoclonal cFos antibodies that target near the N and C terminus together in each IHC to increase the SNR of the cFos signal during imaging. Then, the sections were washed and then incubated in a secondary antibody solution overnight at 4°C (Table 1) and mounted using Fluoromount with DAPI (Sigma). Primary and secondary antibody solutions were made in PBS with 6% normal donkey serum and 0.3% Triton-X. Whole brain images were taken at 10X magnification using an Olympus VS200 Slide Scanner in the Biological Imaging Facility at Northwestern University [RRID: SCR_017767]. The exposure time for each channel was kept consistent across imaging sessions to minimize batch effects of imaging.

**Table 1.** IHC antibodies and concentrations. List of antibodies and tracers used in this study. The dilution of all secondary antibodies and streptavidin was 1:500. All antibodies used in this study have been previously validated by the manufacturers. Validation data and protocols are available on the respective manufacturers’ websites.

| Primary Ab / Tracer | Dilution | Secondary Ab |
| --- | --- | --- |
| Mouse anti-NeuN<br>(Millipore Sigma, MAB377) | 1:500 | Donkey anti-mouse Alexa 488<br>(Abcam, ab150105) |
| Rabbit anti-cFos (9F6)<br>(Cell Signaling, #2250) | 1:500 | Donkey anti-rabbit Alexa 546<br>(Invitrogen, A10040) |
| Rabbit anti-cFos (E2I7R)<br>(Cell Signaling, #31254) |  |  |

### Whole-brain analysis and cFos detection

Coronal brain slice images were aligned to the Allen Atlas Common Coordinate Framework v3 (CCFv3) using Aligning Big Brains and Atlases^47^ (ABBA; v0.10.4) as reported previously. Detection of cFos was performed using StarDist^42^ run through QuPath (v0.6.0)^104^. We describe our pipeline for whole-brain analysis and cFos detection briefly below.

#### ABBA Atlas Registration (Figure S1)

1. First, the slices were placed in order and then flipped to ensure the same orientation. Then, the cryostat cutting angle was set and the slices aligned along the A/P axis.
2. Next, we performed affine registration followed by spline registration two times, comparing DAPI to Nissl, NeuN to Ara, and cFos to Ara. We used 10x resampling for affine registrations and 15x resampling and 20 gridpoints for spline registration.
3. Finally, we manually registered partial sections and manually adjusted registrations as needed.
4. To exclude missing regions or regions with significant damage, we excluded tissue with an average NeuN signal below 25% of the median for a slice from all further analysis.

#### cFos Detection (Figure S2)

1. cFos-positive cells were identified using StarDist run through Qupath using the pretrained *dsb2018_heavyaugment.pb model*. The model is archived on GitHub with the analysis scripts.
2. Detections were then filtered based on size (5 - 22 μm diameter), circularity (> 0.5), and StarDist detection probability (> 0.25).

We wrote a custom groovy script to run the cFos detection and import Allen Atlas CCFv3 registrations from ABBA into QuPath which is available on GitHub. In brief, this script: runs StarDist to detect putative cFos detection; filters putative detections by size, circularity, and StarDist detection probability to get cFos positive cells; imports ABBA CCFv3 registrations; marks cFos positive cell position in CCFv3 space; excludes missing regions based on NeuN average fluorescence.

The coordinates of cFos positive detections in CCFv3 space were exported for each brain. Additionally, cFos positive cell counts for each brain area as well cFos positive cell density, defined as

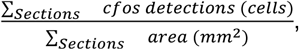

were exported and calculated for each brain.

#### cFos Detection Validation (Figure S2-3)

To validate the cFos detection pipeline, automated detection was compared to expert human counts. Regions throughout the brain including cortex, cerebral nuclei, and brainstem were selected at random sampling from each experimental group. Human counts were made, blinded to experimental groups and detection results. Then, each human and automated count were compared to determine the overlap, finding that generally the cFos detection algorithm overcounts compared to the expert, human experimenter (Figure S3E). In contrast, in regions with dense cFos expression (e.g. cortex or the cerebellar granule layer) the algorithm undercounts, due to the inability to segment individual neurons (shown in Figure S4). Thus, we chose a detection threshold where the undercounting and overcounting were balanced. Notably, most of the algorithm’s apparent false positives — cells it detected that the human counter missed — were confirmed as true cFos+ cells on post-hoc review. This suggests the reported error rates likely overstate the algorithm’s true false-positive rate, since human counts were treated as ground truth without this correction.

Critically, the error is not expected to be the same across regions as neuronal size and density appear to matter, but the error is conserved between each experimental group. Thus, the relative cFos density between regions should not be used definitively. However, within a region the relative differences in cFos expression are accurate and are not expected to change between experimental groups as the IHC, imaging conditions, and cell detection parameters were conserved across groups.

### Statistical analysis

#### Global Linear Mixed Model (LMM)

A linear mixed model (LMM) was fit to the global cFos density. Global cFos was defined as the unweighted mean across brain regions of log-transformed regional cFos density (cells/mm²), computed over the 338 of 360 CCFv3 regions with at least *n* = 1 in all 8 design cells for that region. Taking the unweighted mean across regions ensures that large areas do not disproportionately contribute to the global model. The fixed effects were sex (male and female), genotype (control and MKO), and light (light- and dark-exposed) as well as all interactions. The effects were coded as control, male, and light-exposed as positive and MKO, female, and dark-exposed as negative. Thus, a positive value on the main effect of light means higher cFos in the light than in the dark. Reported effects are reconstructed linear combinations of the fixed effects. Sixteen contrasts were computed as planned decompositions of the three-way interaction: eight simple effects (Figure 2E-F; light within each genotype x sex cell, and genotype within each sex x light cell), four two-way interaction contrasts (Figure 2D; genotype x light and genotype x sex by the third factor), three main effects (Figure 2C), and the omnibus three-way interaction (Figure 2C).

IHC batch (*N* = 6) was fit as a random intercept (variance estimate = 0.072, SE = 0.237; residual scale = 0.406), as significant batch-to-batch variability in global density was observed. Perfusion batches were balanced by light and typically genotype, but were often restricted to one sex. However, perfusion batches were pooled for IHC, permitting an estimate of the random intercept. While single-sex perfusion batches are a limitation, the observed crossover effect cannot be driven by an IHC-batch driven shift of group means uniformly, thus the sign-reversal in the genotype-dependent light response cannot be attributed to batch structure.

The mixed-linear model was implemented in Python 3.13 using statsmodels and MixedLM. The model was fit by restricted maximum likelihood (REML) with the L-BFGS optimizer and converged (log-likelihood = −38.86, 40 observations in 6 groups). Inference on fixed effects and contrasts used Wald tests, reported against both a normal reference and a t reference with df = 32.

We validated the model’s robustness using three strategies (Figure S4). First, to ensure the model was robust to the optimizer used and starting point, we performed the mixed-linear regression while varying the optimizer and starting point. We found that the three-way interaction is significant regardless of the optimizer (L-BFGS, conjugate gradient, Powell, BFGS) and starting point (Figure S4A). Second, to ensure that no individual IHC batch was driving the observed effect, we systematically removed all the samples from one IHC batch and reran the analysis. We found that the three-way interaction is significant even with the removal of each batch (Figure S4B). Finally, to ensure that no individual samples were driving the observed patterns, we systematically removed each sample and reran the analysis, finding that individual animals did not drive the effect (Figure S4C). As a further check on the batch-correction strategy, the model was refit on two alternative constructions of the global scalar with and without IHC batch-residualization: a volume-weighted mean and the root density (total cFos counts per brain / total area sampled per brain) (Figure S4D). The three-way interaction was consistent in sign, magnitude, and significance across all specifications.

#### Per-Region Linear Model

Linear models were fit based on the cFos densities for each brain region, using log-transformed cFos densities after the addition of a pseudocount to account for zeros in the dataset. Each CCFv3 grey matter region (*N* = 360) was fit using an ordinary least squares linear model with log(cFos density) ∼ sex x genotype x light. A per-region linear model was fit for each contrast in which each design cell had at least (*n* = 1). For the full dataset, 338 of 360 regions were analyzed, and 22 regions had insufficient data. For the MKO dataset (Figure 4), 348 of 360 regions were analyzed and 12 regions had insufficient data. Analyses were run using Python 3.13 using the statsmodels package.

The three-way interaction of sex x genotype x light, and all interactions, were computed for the per-region linear model. All model fits converged. This consisted of three main effects, the three-way interaction, three pooled and six split two-way interactions, and twelve simple effects. Each contrast was specified as a weight vector w (1 x 8), assigning a value to each of the eight design cells. Because the model’s fitted coefficients represent main effects and interactions rather than the eight cells directly, w was converted into that coefficient space via c = wᵀM, where M (8 design cells x 8 coefficients) encodes each cell in terms of the model’s coefficients. The resulting contrast vector c (1 x 8) was then used to calculate the contrast estimate as cᵀβ, where β are the model’s fitted coefficients. Standard errors were calculated as √(cᵀΣ_β_c), where Σ_β_ is the coefficient covariance matrix, and *p*-values are two-sided *z*-tests. Estimates are on a Log_2_ scale. Correction for multiple comparisons was made by Benjamini-Hochberg correction applied across regions within each contrast, to generate corrected *q*-values.

Per-contrast *p* value histograms, standard error distributions, and quantile-quantile plots were inspected prior to interpretation. Further, residual variance did not differ by sex, either pooled across regions (Levene’s test, *p* = 0.33; variance ratio = 0.93) or region by region (6.1% of 342 regions at *p* < 0.05; 5% expected by null; median variance ratio = 0.98), giving no evidence of differences in variance amongst the sexes and supporting pooled models across sex. Regions with near-degenerate variance ratios were flagged, and should be interpreted with caution (N = 4; nucleus ambiguous, ventral (AMBv), linear nucleus of the medulla (LIN), lateral reticular nucleus, parvocellular part (LRNp), spinal nucleus of the trigeminal, interpolar part (SPVI)). Of note, per-animal brainwide density strongly predicts the per-region estimate, and thus, per-region analyses should be considered in context with the brainwide result.

Permutation testing followed the approach from Winkler et al. (2014), on the subset of regions with complete data for every animal (*n* = 271). In brief, interaction terms were approximated by Freedman-Lane permutation testing of model residuals^105^, as no permutation of design variables can null an interaction term while leaving the real main effects and lower-order interactions intact, whereas simple effects and main effects were tested with permutation of design labels. As light is the only experimentally randomized factor, it was permuted where possible. For contrasts where light is held constant (eg. Figure 4), sex or genotype were permuted instead. 2000 permutations were drawn per contrast, and P values were computed as (1 + k) / (*N* +1), where k is the number of permutations at least as extreme as the observed and *N* is the number of permutations^106^. Thus, for *N* = 2000 permutations, the *p* value floor would be 1/2001 ≈ 0.0005. Where the number of distinct labellings available to a restricted permutation was smaller than this, that resolution ceiling is reported instead, as no additional permutations can lower it.

The breadth of the brainwide effect was measured as the mean *z* across regions (two-sided). Enrichment was tested as mean *z*_subset_ - mean *z*_all_. Sets were defined based on prior literature. Size and contiguity matched region sets were generated (*N* = 5000) as the null for enrichment testing. Monte Carlo standard errors of the permutation *p* are reported.

The sets for enrichment tests were the following: retinorecipient – anterior hypothalamic nucleus (AHN), bed nucleus of the stria terminalis (BST), dorsal terminal nucleus of the accessory optic tract (DT), intergeniculate nucleus (IGL), LGd, LGv, lateral habenula (LH), lateral peduncle (LP), lateral terminal nucleus of the accessory optic tract (LT), medial amygdala (MEA), nucleus of the optic tract (NOT), olivary pretectal nucleus (OP), periaqueductal grey area (PAG), subparaventricular zone (SBPV), SCN, superior colliculus, optic layer (SCop), suprageniculate nucleus (SGN), substantia innominata (SI), SO, VLPO, zona incerta (ZI), anterior amygdalar area (AAA), anterodorsal nucleus (AD), anterior pretectal nucleus (APN), central lateral nucleus of the thalamus (CL), inferior colliculus, dorsal region (ICd), DR, lateral hypothalamic area (LHA), ventral posterolateral nucleus of the thalamus (VPL), medial pretectal area (MPT), midbrain reticular nucleus (MRN), medial terminal nucleus of of the accessory optic tract (MT), parabrachial nucleus (PB), paranigral nucleus (PN), peripuduncular nucleus (PP), posterior pretectal nucleus (PPT), retrochiasmatic area (RCH), subgeniculate nucleus (SubG), magnocellular nucleus (MA), diagonal band nucleus (NDB)^33,34^; ipRGC targets – AHN, BST, DT, IGL, LGd, LGv, LH, LP, LT, MEA, NOT, OP, PAG, SBPV, SCN, SCop, SGN, SI, SO, VLPO, and ZI^35–37^; image-forming region – VISal, VISam, VISl, VISp, VISpl, VISpm, VISli, VISpor, VISa, VISrl, TEa, RSPagl, RSPd, RSPv^51,52^; neuromodulatory – NDB, SI, MA, medial septal nucleus (MS), SO, PVH, DR, CLI, LC, SNc, VTA, PPN, and LDT, acetylcholine system – NDB, SI, MA, MS, PPN, and LDT^62–64^, dopamine system – SNc and VTA^66,67^, and oxytocin system – SO and PVH^65^.

#### Contrast Task Partial Least Squares Correlation (PLSC)

We performed contrast task PLSC^61^ to identify patterns in regional cFos covariance throughout the brain that is attributable to each experimental contrast. Where the per-region linear models define which individual regions vary between groups, PLSC measures spatial patterns of brain regions that change between groups.

PLSC operated on the 338 brain regions with data in each design-cell (i.e. group) using means rather than on individual animals, with cells weighted equally, which keeps the contrasts exactly orthogonal despite unequal cell N, ensuring unequal sample size does not bias the weights. Regional densities were log-transformed after addition of a pseudocount equal to half the smallest non-zero density. IHC batch was then removed by fitting a per-region linear mixed model (log density ∼ genotype x sex x light, batch as random intercept) and subtracting only the batch-attributable shift (BLUP), leaving the fixed effects intact. Since PLSC has no random effect structure, this is necessary to distinguish IHC batch effects from a design effect. Additionally, given the brainwide impacts on gain, this global signal would be the largest source of the variance in the dataset unless removed. Thus, each animal’s brainwide mean was subtracted from each region, and the latent variables defined by this analysis therefore represent spatial redistribution relative to the animal’s mean. Data was then normalized within brain regions by *z*-scoring across all animals so that individual regions do not dominate the decomposition based on scale alone.

The design was coded as a ±1 contrast matrix over the eight groups (with males, control, and light +1), with columns for three main effects, three two-way interactions, and the three-way interaction. The cross-covariance matrix R = KᵀL was calculated, where K is the 8 x 7 matrix of contrasts and L is the 8 x 338 matrix of design-cell means for each region. Each row of R is the covariance of one contrast across regions. The magnitude of each contrast was calculated as the norm of its row of R across regions, giving a multivariate magnitude of that contrast. Because the contrasts are orthogonal and identically scaled, contrast magnitudes are comparable across contrasts. The brain salience for each region was calculated by row normalization to unit length, putting the brain saliences on a common scale.

For comparison, we also conducted task-centered PLSC as described in Chiaruttini et al., 2025, by decomposing the grand-mean-centered design-cell matrix by singular value decomposition, yielding 7 latent variables (LVs) with design saliences (how much each contrast contributes), singular values (covariance captured), and brain saliences (per-region contribution) for each of the 338 tested brain regions. Per-region claims for significance are made based on the corrected *q*-value from the per-region linear model run using PLSC normalized data in which per-animal mean was subtracted. The reliability of each latent variable was determined by resampling by splitting over perfusion batches (5,000 splits). The correlation between halves is reported, and we report the overlap of the top 25 regions based on absolute salience (chance = 0.048 Jaccard overlap). No LV showed strongly reliable patterns, and thus the LVs should be interpreted cautiously.

Contrast and LV significance was assessed by 10000 permutations, comparing each observed contrast magnitude or LV singular value against its null distribution (one-tailed, because both are non-negative norms)., and *p* values were computed as (1 + k) / (*N* +1), where k is the number of permutations at least as extreme as the observed and *N* is the number of permutations^106^. Two permutation schemes were based on the ability to generate null populations for the dominant design salience. Light and genotype were randomized in their perfusion batches so permutations were within batch. Sex was not assigned, and thus, sex was permuted freely across animals, so tests involving sex are population-level tests of association rather than randomization tests. *p* values between groups are not comparable. Perfusion batch permutation was used for LVs, the main effects of light and genotype, and the two-way effect of light x genotype. The sex permutation was used for the main effect of sex, the two-way effects of genotype x sex and light x sex, as well as the three-way interaction. Of note, this creates a null for the sex associated structure of the interaction contrasts, but it can not fully create a null for the interaction.

Region-level contributions were assessed by bootstrap resampling of the (unnormalized) cross-covariance matrix, stratified within design cell (10,000 resamples). No rotation is needed here since the contrast weights are fixed rather than estimated by decomposition. Bootstrap ratios (BSR) were calculated as the mean bootstrapped value divided by its bootstrap standard error, and regions with |BSR| > 2 are reported as stable contributors. Note, we do not use BSR as a significance measure and instead consider it a stability measure.

### Data visualization

Data was visualized using the BrainGlobe^107^ package in Python 3.13 using custom scripts (http://github.com/schmidtlab-northwestern) and GraphPad Prism 10.6.1. Heatmaps show mean for the entire region.

## RESOURCE AVAILABILITY

### Data Availability

Raw imaging data is available upon request. Compiled data is available as cFos counts and density for each brain region in each mouse as well as coordinates of each cFos positive cell in the Allen Atlas CCFv3 on Github (http://github.com/schmidtlab-northwestern). Data is organized by brain with filenames of MouseID_annotation_measurements.csv (cFos counts, region area, and cFos density for each region of each section) and MouseID_detection_measurements.csv (Coordinates of each cFos detection in Allen Atlas CCFv3 Mouse Brain Atlas). A metadata file, titled mouse_info.csv, is available and includes sex, genotype, perfusion batch, IHC batch, age, and light conditions for each mouse. Additionally, a compiled file of cFos density / regions for each mouse in each condition is available, titled density_by_region.csv. This data can be explored interactively at [website].

### Code Availability

The code for data analysis and visualization is available on GitHub (http://github.com/schmidtlab-northwestern).

### Lead Contact

Further information and requests for resources and reagents should be directed to and will be fulfilled by the lead contact, Dr. Tiffany Schmidt.

## ACKNOWLEDGEMENTS

This work was funded by National Institutes of Health grant DP2 EY027983 (TMS), National Institutes of Health grant T32 EY025202, National Institutes of Health grant F31 EY036265 (JDB), and the Christina Enroth-Cugell Graduate Research Award (JDB).

We thank Dr. Yevgenia Kozorovitskiy and Sara Freda for their assistance with slide scanner imaging and their advice and expertise regarding development of whole-brain atlases. We also thank Dr. Tong Zhang at the Biological Imaging Facility at Northwestern University for working with us to procure the Olympus VS200 Slide Scanner. We also thank Dr. Emma Alexander for advice on developing the cFos detection pipeline. Finally, we thank all of the other members of the Schmidt Lab for their help and support on this project throughout the years!

## AUTHOR CONTRIBUTIONS

Conceptualization: JDB and TMS. Methodology: JDB and MRS. Investigation: JDB, MRS, ACW, AK, LG, and TMS. Visualization: JDB, MRS, and TMS. Funding acquisition: JDB and TMS. Project administration: TMS. Supervision: TMS. Writing – original draft: JDB and TMS. Writing – review and editing: JDB, MRS, AK, LG, ACW, and TMS.

## SUPPLEMENTAL TABLES AND FIGURES

**Figure S1.**
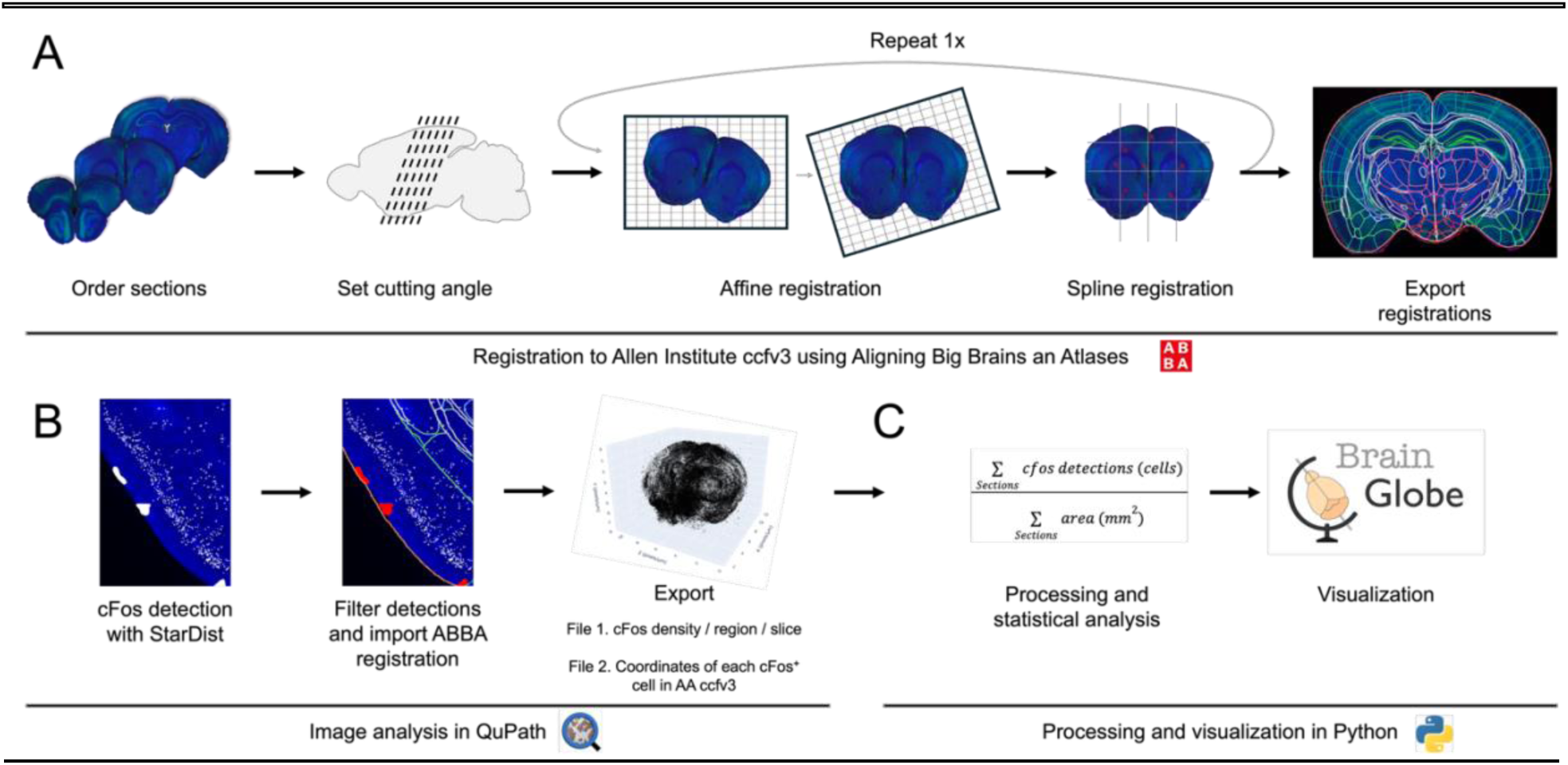
Whole-brain analysis pipeline. (A) Registration to the Allen Mouse Brain Reference Atlas (CCFv3) using ABBA. (B) cFos detection, filtering and registration import in QuPath, and data export in Qupath. (C) Post-processing and visualization using Python.

**Figure S2.**
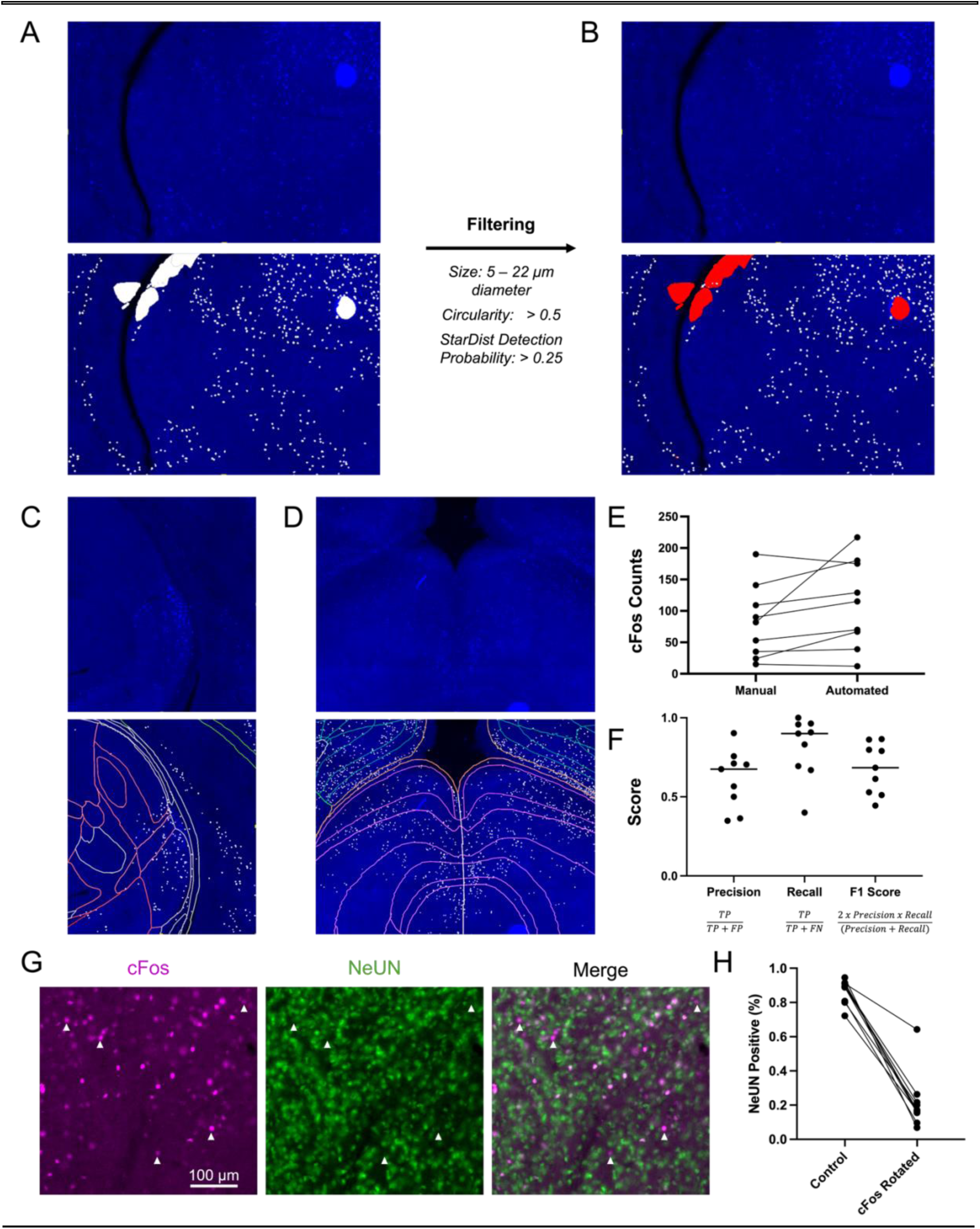
cFos detection pipeline and validation. (A) cFos detection using StarDist. (B) Filter cFos detections based on size (5 - 22 μm diameter), circularity (circularity > 0.5), and StarDist detection probability (detection probability > 0.25). (C) Example cFos detection and registration in the lateral geniculate nucleus of a control, light exposed female. (D) Example cFos detection and registration in the superior colliculus of a control, light exposed female. (E) Counts compared to expert, human quantifiers. (F) Validation of cFos detection across key metrics. Abbreviations – TP: true positive; FP: false positive; FN: false negative. (G) Representative image of cFos (Magenta) and NeuN (Green). (H) Percent of cFos+ cells that colocalize with NeuN in unrotated control images as well as images with the cFos channel rotated 180°.

**Figure S3.**
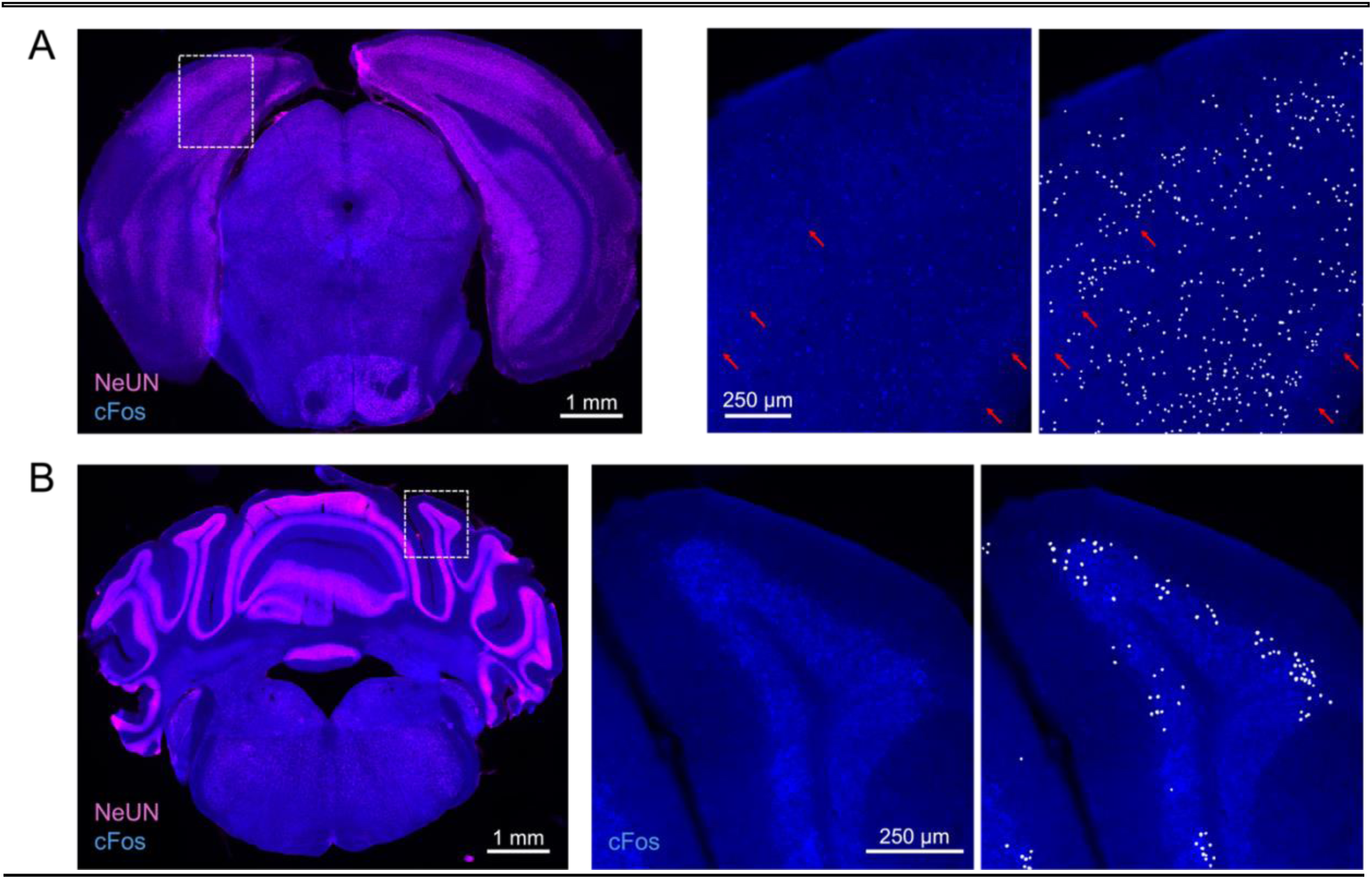
cFos detection undercounting is more severe in cortex and cerebellum. (A) cFos detection in the isocortex. Red arrows denote missed cFos detections. (B) cFos detection in the cerebellum.

**Table S1.** Global LMM results.

| Contrast | Estimate ($\text{Log}_2$) | Standard Error ($\text{Log}_2$) | p value |
| --- | --- | --- | --- |
| **Light | 0.92 | 0.30 | 0.0025 |
| *Genotype | -0.78 | 0.35 | 0.0241 |
| Sex | -0.07 | 0.33 | 0.8292 |
| **Sex x Genotype x Light | 3.78 | 1.24 | 0.0024 |

**Table S2.** Planned decompositions of global LMM results.

| Contrast | Estimate (Log<sub>2</sub>) | Standard Error (Log<sub>2</sub>) | p value |
| --- | --- | --- | --- |
| Genotype x Light \| M | 1.30 | 0.81 | 0.1105 |
| **Genotype x Light \| F | -2.48 | 0.95 | 0.0090 |
| Sex x Light \| Control | 1.37 | 0.84 | 0.1016 |
| **Sex x Light \| MKO | -2.41 | 0.91 | 0.0077 |
| *Light \| Control, M | 1.31 | 0.62 | 0.0346 |
| Light \| Control, F | -0.06 | 0.58 | 0.9167 |
| Light \| MKO, M | 0.01 | 0.53 | 0.9717 |
| ***Light \| MKO, F | 2.42 | 0.72 | 0.0008 |

**Figure S4.**
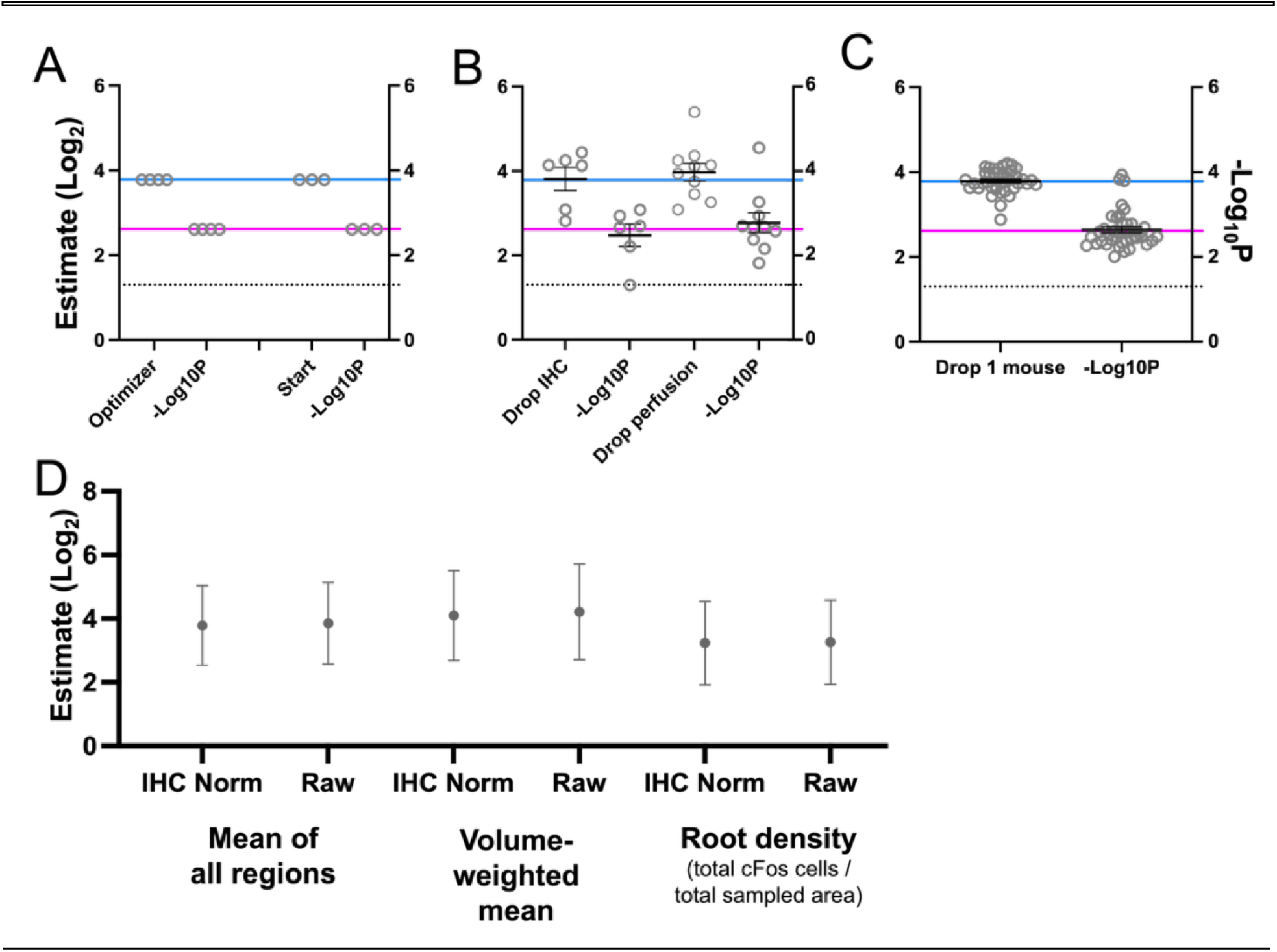
Brainwide cFos expression mixed linear model robustness. (A) The model converges and has a nearly identical estimate and *p*-value using each tested optimizer (*N* = 4) and random starts (*N* = 3). (B) Three-way interaction holds when systematically removing each individual mouse. Data are Mean ± SEM. (C) Three-way interaction holds when systematically removing each IHC and perfusion batch. Data are Mean ± SEM. (D) Three-way interaction holds when using different methods for calculating global density with and without IHC batch removed. Solid lines designate the estimate (blue) and *p*-value (pink) of the model in Figure 2 (IHC normalized mean of all regions). Dotted line designates *p* = 0.05.

**Figure S5.**
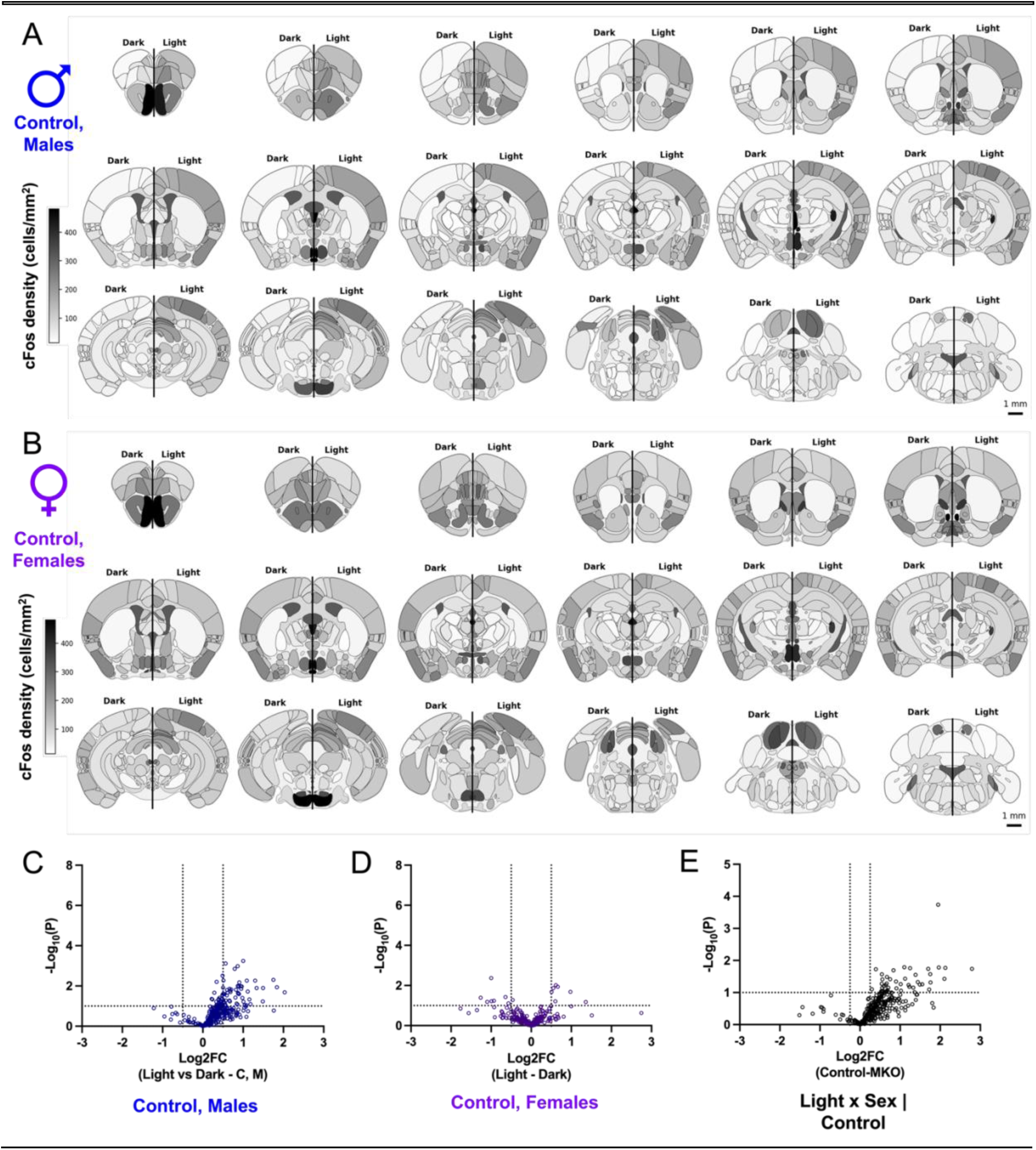
Atlas of light induced cFos expression. (A) Heatmaps of whole-brain cFos density in male dark-(left) and light-exposed controls (right) (mean; *n* = 4 light and 5 dark). (B) Heatmaps of whole-brain cFos density in female dark-(left) and light-exposed controls (right) (mean; *n* = 5 light and 6 dark). (C) Volcano plot of the Log_2_FC of the effect of light in control males and the uncorrected -Log_10_(*p*) for each brain region. (D) Volcano plot of the Log_2_FC of the effect of light in control females and the uncorrected -Log_10_(*p*) for each brain region. (E) Volcano plot of the Log_2_FC of the interaction of sex x light in controls and the uncorrected -Log_10_(*p*) for each brain region. Heatmaps are every 0.5 mm along the A/P axis of the brain. Scale bar = 1 mm.

**Figure S6.**
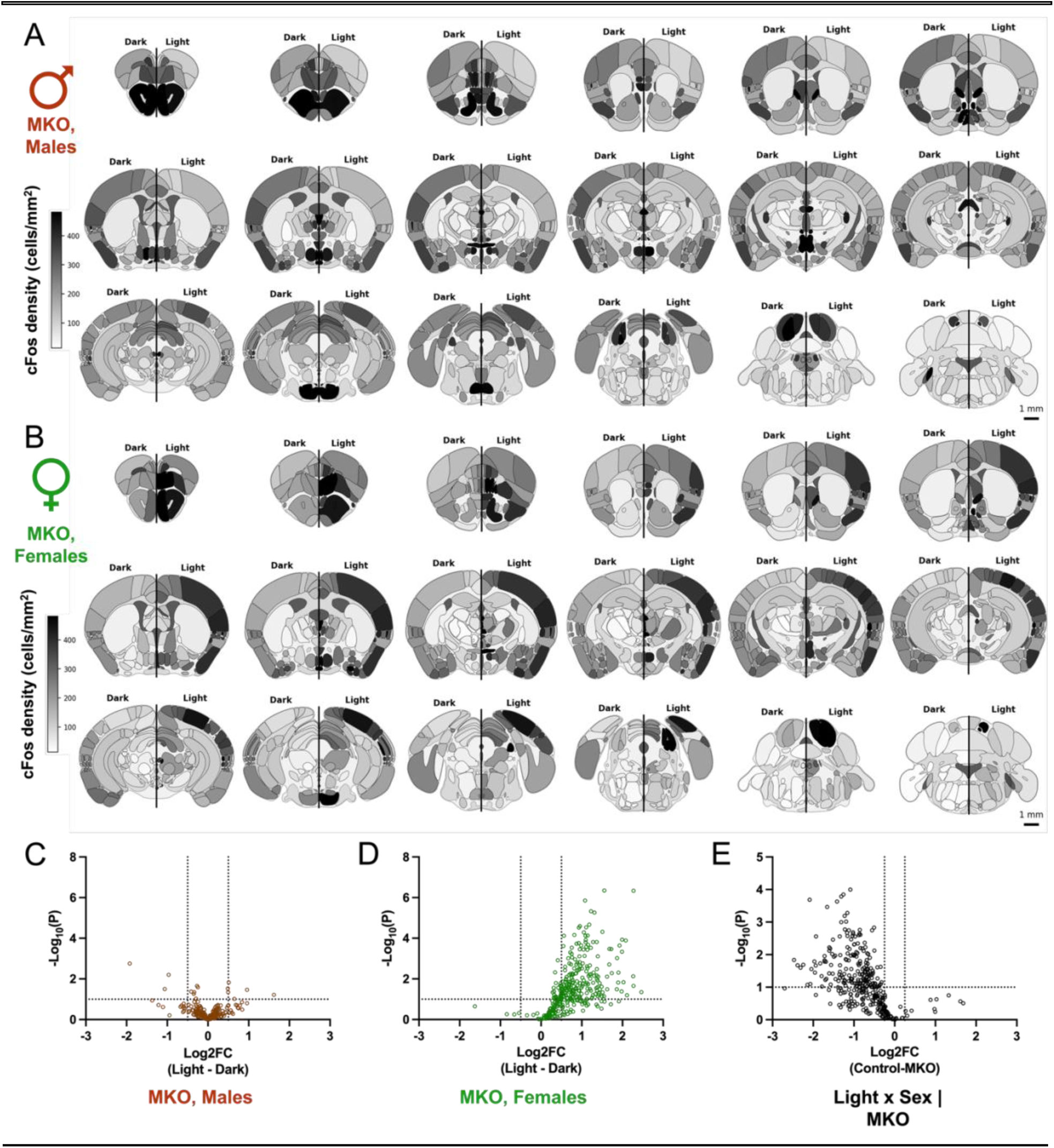
Light induced cFos expression is attenuated in MKO males but unmasked in MKO females. (A) Heatmaps of whole-brain cFos density in male dark-(left) and light-exposed MKOs (right) (mean; *n* = 5 light and 8 dark). (B) Heatmaps of whole-brain cFos density in female dark-(left) and light-exposed MKOs (right) (mean; *n* = 3 light and 4 dark). (C) Volcano plot of the Log_2_FC of the effect of light in MKO males and the uncorrected -Log_10_(*p*) for each brain region. (D) Volcano plot of the Log_2_FC of the effect of light in MKO females and the uncorrected -Log_10_(*p*) for each brain region. (E) Volcano plot of the Log_2_FC of the interaction of sex x light in MKOs and the uncorrected -Log_10_(*p*) for each brain region. Heatmaps are every 0.5 mm along the A/P axis of the brain. Scale bar = 1 mm.

**Figure S7.**
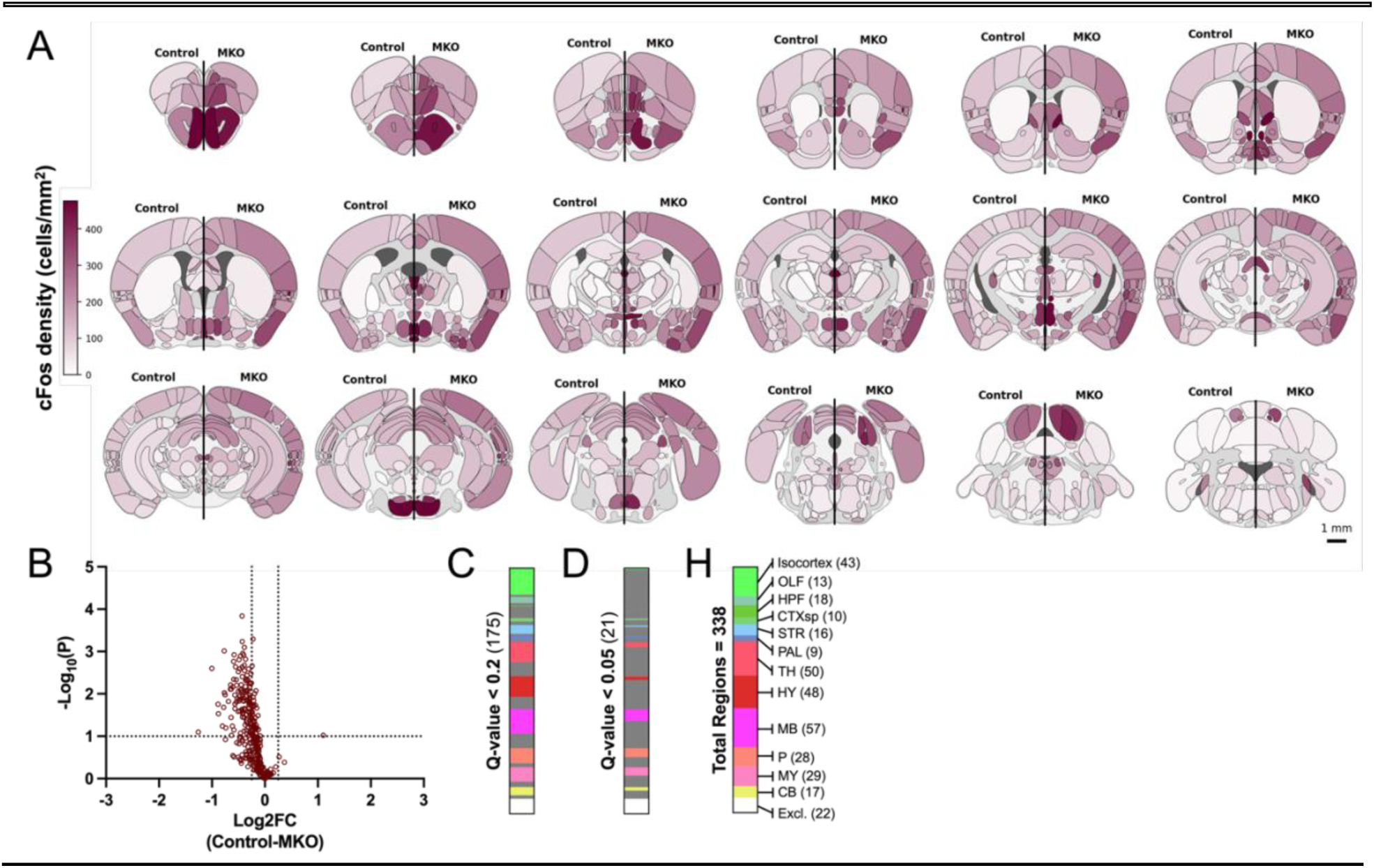
Main effect of genotype. (A) Heatmaps of whole-brain cFos density in controls (left) and MKOs (right) (mean; *n* = 20 controls and 20 MKOs). (B) Volcano plot of the Log_2_FC of the main effect of genotype and the uncorrected -Log_10_(*p*) for each brain region. (C-E) Count of brain regions with a corrected *q* < 0.2 (C) and *q* < 0.05 (D) of the total regions tested (E) in the per region linear model. Heatmaps are every 0.5 mm along the A/P axis of the brain. Scale bar = 1 mm. All data in this figure pools male and female data.

**Figure S8.**
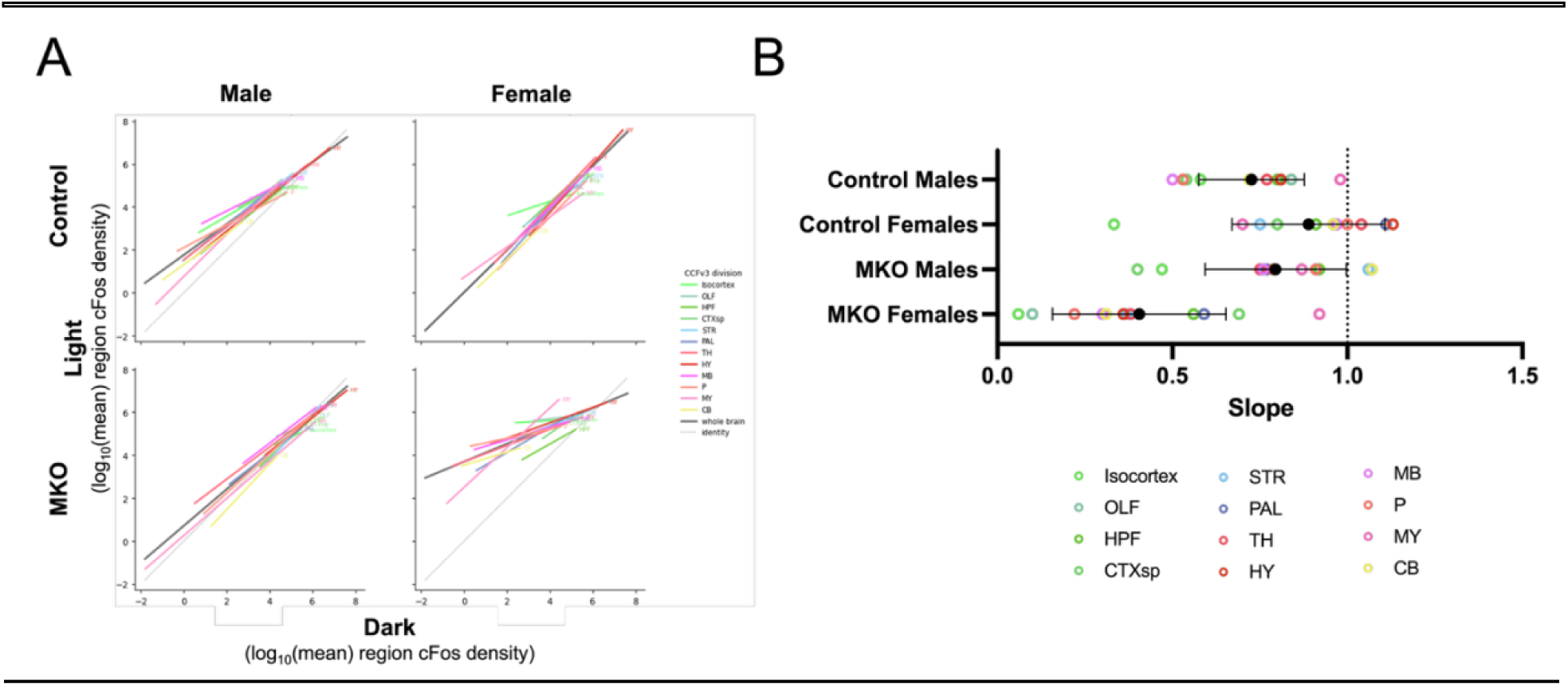
Population light-dependent scaling by subdivision. (A) Fit for each major subdivision of the brain. (B) Slope for the fit of each subdivision compared to the overall mean and standard deviation (black; mean slope ± SE)

**Figure S9.**
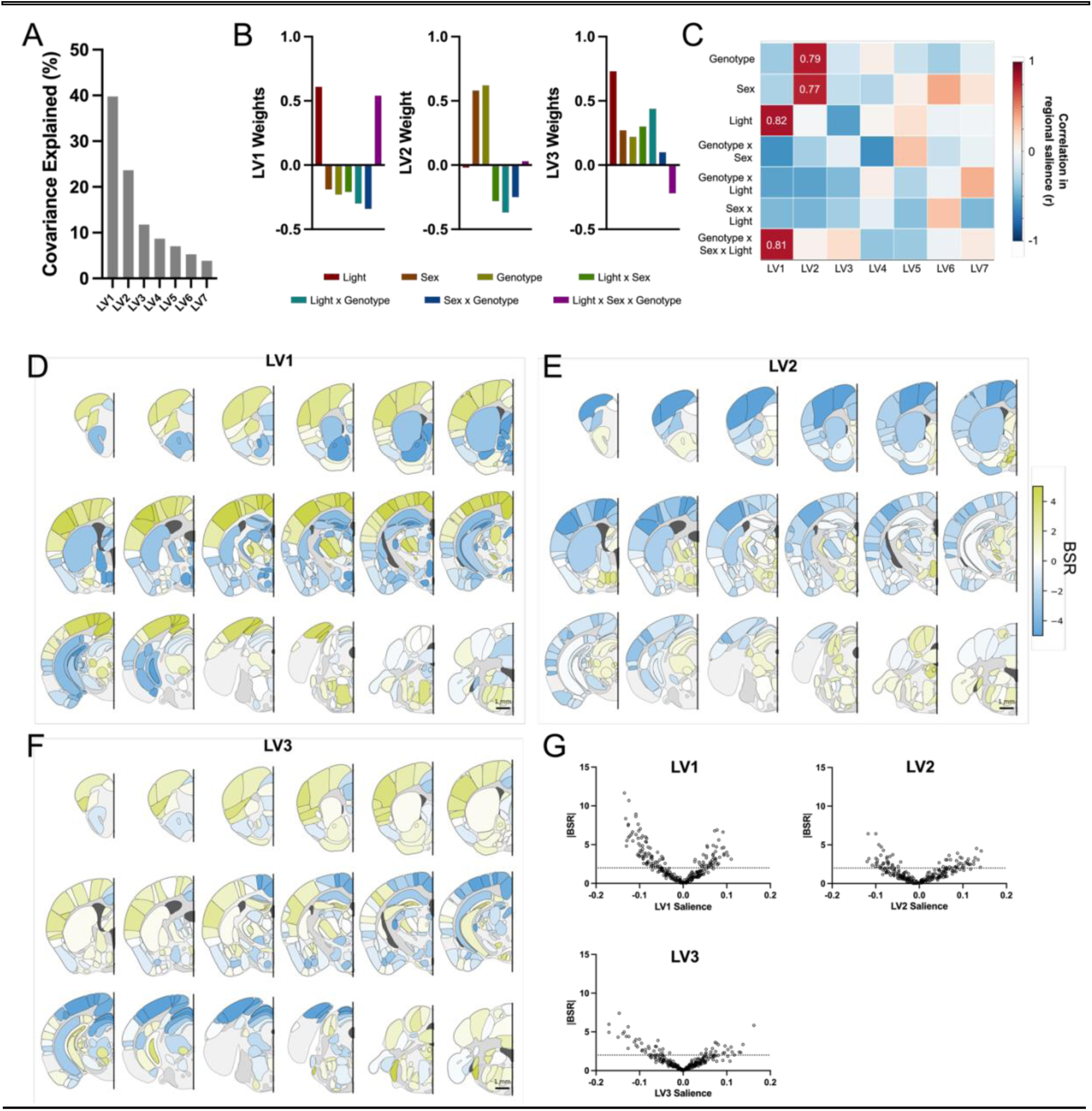
Task-centered PLSC. (A) Covariance explained by each latent variable. (B) Contribution of each contrast to LV1-3. (C) Covariance between per-region saliences of the unrotated contrast (Figure 5) with each LV identified using singular value decomposition. (D-F) Heatmaps of BSR for LV1 (D), LV2 (E), and LV3 (F). (G) Volcano plots of salience and |BSR| for LV1-3. Heatmaps are every 0.5 mm along the A/P axis of the brain. Scale bar = 1 mm.

